# A systems biology approach uncovers the core gene regulatory network governing iridophore fate choice from the neural crest

**DOI:** 10.1101/318402

**Authors:** K. Petratou, T. Subkhankulova, J. A. Lister, A. Rocco, H. Schwetlick, R. N. Kelsh

**Affiliations:** Department of Biology and Biochemistry and Centre for Regenerative Medicine, University of Bath, Bath BA2 7AY, United Kingdom.; Department of Human and Molecular Genetics and Massey Cancer Center, Virginia Commonwealth University School of Medicine, PO Box 980033, Richmond, Virginia 23298, USA.; Department of Microbial and Cellular Sciences, Faculty of Health and Medical Sciences, University of Surrey, Guildford GU2 7XH, UK.; Department of Mathematical Sciences, University of Bath, Bath BA2 7AY, United Kingdom.

## Abstract

Multipotent neural crest (NC) progenitors generate an astonishing array of derivatives, including neuronal, skeletal components and pigment cells (chromatophores), but the molecular mechanisms allowing balanced selection of each fate remain unknown. In zebrafish, melanocytes, iridophores and xanthophores, the three chromatophore lineages, are thought to share progenitors and so lend themselves to investigating the complex gene regulatory networks (GRNs) underlying fate segregation of NC progenitors. Although the core GRN governing melanocyte specification has been previously established, those guiding iridophore and xanthophore development remain elusive. Here we focus on the iridophore GRN, where mutant phenotypes identify the transcription factors Sox10, Tfec and Mitfa and the receptor tyrosine kinase, Ltk, as key players. We present expression data, as well as loss and gain of function results, guiding the derivation of an initial iridophore specification GRN. Moreover, we use an iterative process of mathematical modelling, supplemented with a novel, Monte Carlo screening algorithm suited to the qualitative nature of the experimental data, to allow for rigorous predictive exploration of the GRN dynamics. Predictions were experimentally evaluated and testable hypotheses were derived to construct an improved version of the GRN, which we showed produced outputs consistent with experimentally observed gene expression dynamics. Our study reveals multiple important regulatory features, notably a *sox10*-dependent positive feedback loop between *tfec* and *ltk* driving iridophore specification; the molecular basis of *sox10* maintenance throughout iridophore development; and the cooperation between *sox10* and *tfec* in driving expression of *pnp4a*, a key differentiation gene. We also assess a candidate repressor of *mitfa*, a melanocyte-specific target of *sox10*. Surprisingly, our data challenge the reported role of Foxd3, an established *mitfa* repressor, in iridophore regulation. Our study builds upon our previous systems biology approach, by incorporating physiologically-relevant parameter values and rigorous evaluation of parameter values within a qualitative data framework, to establish for the first time the core GRN guiding specification of the iridophore lineage.

**Author Summary:** Multipotent neural crest (NC) progenitors generate an astonishing array of derivatives, including neuronal, skeletal components and pigment cells, but the molecular mechanisms allowing balanced selection of each fate remain unknown. In zebrafish, melanocytes, iridophores and xanthophores, the three chromatophore lineages, are thought to share progenitors and so lend themselves to investigating the complex gene regulatory networks (GRNs) underlying fate segregation of NC progenitors. Although the core GRN governing melanocyte specification has been previously established, those guiding iridophore and xanthophore development remain elusive. Here we present expression data, as well as loss and gain of function results, guiding the derivation of a core iridophore specification GRN. Moreover, we use a process of mathematical modelling and rigorous computational exploration of the GRN to predict gene expression dynamics, assessing them by criteria suited to the qualitative nature of our current understanding of iridophore development. Predictions were experimentally evaluated and testable hypotheses were derived to construct an improved version of the GRN, which we showed produced outputs consistent with experimentally observed gene expression dynamics. The core iridophore GRN defined here is a key stepping stone towards exploring how chromatophores fate decisions are made in multipotent NC progenitors.

## Introduction

Despite decades of work, we still have only a superficial idea of how stem cells generate their distinct derivatives. This question becomes more acute if we consider that these fate choices are often made in a physically constrained environment (e.g. a stem cell niche), suggesting that fate-specification by environmental signals may be only part of the mechanism.

Neural crest cells (NCCs) are a multipotent embryonic cell-type, sharing many properties with stem cells and indeed being retained as adult neural crest stem cells in various niches (Shakhova and Sommer 2008). They are an important model for understanding the genetics of stem cell fate choice, since they generate a fascinating diversity of derivative cell-types, including many peripheral neurons, all peripheral glia, various skeletogenic cells, and pigment cells (Gilbert 2000; Le Douarin and Dupin 2003; Trainor 2014). The latter are restricted to melanocytes in mammals, but are much more diverse in the anamniotes, such as fish (Kelsh 2004; Bagnara 2007; Schartl et al. 2016). In the well-studied zebrafish, there are three distinct types of pigment cells, namely black melanocytes, iridescent iridophores and yellow xanthophores, and in medaka, these three are supplemented by white leucophores. It is a long-standing, although largely untested, proposal that all pigment cells (or chromatophores) share a common origin from a neural crest (NC) derived, partially-restricted pigment cell progenitor, a chromatoblast (Bagnara et al. 1979; Lopes et al. 2008). This, in conjunction with the inherent genetic tractability of these cell types, makes study of pigment cell development from the NC an exciting ‘model within a model’ for the genetics underlying stem cell fate choice.

It is generally assumed that NC fate specification follows a progressive fate restriction model, with early, fully multipotent NCCs giving rise to individual pigment cell fates via a series of partially-restricted intermediates, and with fate choice consisting of a series of binary choices until a single fate is adopted (Weston 1991; Le Douarin, Calloni, and Dupin 2008). This view is crystallised in the iconic Waddington landscape model of stem cell development (Waddington 1957). Consistent with this view, aside from the chromatoblast, these partially-restricted intermediates for pigment cells have been suggested to include bipotent Schwann cell precursors, capable of forming melanocytes as well as Schwann cells (Adameyko et al. 2009; Adameyko and Lallemend 2010); bipotent melanoiridoblasts (Curran et al. 2010), and bipotent xantholeucoblasts (Nagao et al. 2014, 2018).

Underpinning the observed fate choices are gene regulatory networks (GRNs) with the emergent property of distinct, stable states of gene expression, each corresponding to the molecular signature of a specific derivative cell-type. To understand stem cell fate choice, it is crucial to identify the key components of these GRNs and their regulatory logic. For pigment cell development, genetics has identified a small set of genes crucial for the control of lineage specification and differentiation (Kelsh et al. 1996; Lister et al. 1999; Kelsh 2004; Minchin and Hughes 2008). Integrating studies of these key mutants focused on identifying the core melanocyte GRN. Melanocyte specification centres on expression and maintenance of Microphthalmia-related transcription factor a (Mitfa), a bHLH-Leu Zipper transcription factor that functions as a master regulator of melanocyte development (Lister et al. 1999; Elworthy et al. 2003; Greenhill et al. 2011). Initial expression of *mitfa* depends upon the Sry-related HMG-box 10 (Sox10), a transcription factor shown to directly regulate *mitfa* expression, cooperating with Wnt signalling (Potterf et al. 2000; Elworthy et al. 2003; Vibert et al. 2017). Sox10 plays a similar role in specification of both xanthophores and iridophores as well (Kelsh and Eisen, 2000;Dutton et al. 2001). In the case of iridophores, as well as Sox10, the receptor tyrosine kinase Leukocyte tyrosine kinase (Ltk) plays a crucial role, with loss of function mutants lacking embryonic and adult iridophores, and constitutively activated Ltk signalling driving NCCs to adopt an iridophore fate (Lopes et al. 2008; Rodrigues et al. 2012; Fadeev et al. 2016). We have shown that *ltk* expression appears to show two phases, one in early NC development which we propose represents a multipotent, chromatoblast-like progenitor, and a second in the definitive iridophore lineage (Lopes et al. 2008). Importantly, *mitfa* mutants show an intriguing increase in iridophores accompanying the absence of melanocytes, suggesting a close relationship between these two fates, and interpreted as revealing a shared bipotent progenitor, a melanoiridoblast (Lister et al. 1999; Curran et al. 2010). Mitfa belongs to a subfamily of related transcription factors containing the Transcription Factor E factors; one of these, *tfec*, is expressed in early NCCs, but later in a pattern strikingly reminiscent of iridophores, and is a strong candidate for a master regulator of iridophore development (Lister et al. 2011, Petratou et al., in prep). Finally, the Forkhead box D3 transcription factor (Foxd3) has been proposed to repress *mitfa* expression, consistent with the suggested role of FOXD3 in repressing MITF in other models (Kos et al. 2001; Thomas and Erickson 2009), thus biasing bipotent melanoiridoblast progenitors towards an iridophore fate (Curran, Raible, and Lister 2009; Curran et al. 2010). Although endothelin receptor Ba (Ednrba) shows an expression pattern that marks iridophore development, *ednrba* mutants show no discernible embryonic iridophore phenotype, although they do show loss of iridophores in adults (Parichy et al. 2010). Finally, *pnp4a* has been identified as a useful differentiation marker for the iridophore lineage (Curran et al. 2010).

However, these key genetic insights have yet to be integrated into a comprehensive GRN of pigment cell progenitors, the analysis of which might lead to understanding of how the NC generates each cell-type, and in appropriate ratios. As a first step in this, we have identified a core GRN for melanocyte fate specification (Greenhill et al. 2011). As the number of components of a GRN increase, the standard network diagrams used to depict them become increasingly difficult to interpret using intuition alone. Importantly, therefore, we used an iterative cycle of experimental observations and mathematical modelling to more rigorously assess the GRN as we developed this core model. Using a similar approach, we have subsequently integrated the biphasic role of Wnt signaling in melanocyte development (Vibert et al., 2016).

As a next step towards developing an integrated GRN for pigment cell fate-specification in zebrafish, we here extend the combined use of experimental genetics and mathematical modelling to develop a core GRN for iridophore specification. Many of the experimentally identified key genes in iridophore development, specifically *ltk*, *tfec*, *sox10* and *mitfa,* show multiphasic expression in the NC and so we begin by outlining a working definition of the phases of iridophore specification from early, fully multipotent NCCs to differentiated iridophores. We then use this framework to allow careful interpretation of the highly dynamic gene expression patterns of the key genes in both wild-type (WT) and appropriate mutant embryos to assess the regulatory relationships between them. We refine the mathematical modelling approach developed to analyse the melanocyte GRN (Greenhill et al. 2011), using a literature search to limit parameter space to a reasonable physiological range, and Monte Carlo simulations to assess the robust predictions of GRN models throughout that parameter space. We emphasize that our Monte Carlo approach is particularly suitable in all those cases when quantitative data are not available, but rather qualitative behaviours are known. Supplemented by this approach for model selection, we then use our systems biology framework as a tool to rigorously evaluate a set of related models, refining and expanding them to define the first core GRN for iridophore development in zebrafish.

## Results

### Identification of the iridophore lineage throughout zebrafish embryogenesis

To interpret the expression dynamics of genes of interest during iridophore development in wild-type (WT) embryos, as well as changes of expression patterns in different loss of function contexts, it was crucial to distinguish cell populations comprising the different stages of iridophore development. Gene expression in the zebrafish NC is highly dynamic, reflecting both the rapid fate specification and differentiation of NC derived lineages in zebrafish and the multiphasic expression patterns of many key genes. For example, in previous studies of the *ltk* marker, we have proposed at least three phases of expression, one in multipotent premigratory progenitors, and two representing iridoblasts and differentiated iridophores respectively (Lopes et al. 2008). Tfec was first identified as a Mitf-related bHLH-ZIP transcription factor expressed in premigratory NC and later in a pattern reminiscent of iridophores (Lister et al. 2011). We have recently shown that Tfec is crucial for fate specification of the iridophore lineage, and that *tfec* expression labels both early iridoblasts and differentiated iridophores (Petratou et al, in prep.). Building on these previous studies, we assessed the spatio-temporal locations of presumed iridophore progenitors at key stages of embryogenesis by examining *tfec* expression in whole-mount embryos. WISH on single embryos at 72 hpf confirmed that *tfec* is a definitive marker of differentiated iridophores (Fig 1A), similar to the established iridophore lineage marker, *ltk* (Lopes et al. 2008). Moreover, double labelling of *ltk* and *tfec* expression using multiplexed fluorescent RNAscope revealed that *tfec* is expressed throughout iridophore development (Fig 1C). Furthermore, *tfec* transcripts were first seen in premigratory NC at very early stages, considerably before *ltk* (Fig 1B). Consequently, we interpret *tfec* expression as an excellent marker of ‘iridophore potential’ during zebrafish embryogenesis and use it here to produce a working classification of the stages of iridophore development. We note explicitly that this classification is intended to provide a framework for interpretation of mutant phenotypes, and that assessment of *tfec* alone cannot provide insight into the multipotency of cells at any specific stage.

**Figure 1.**
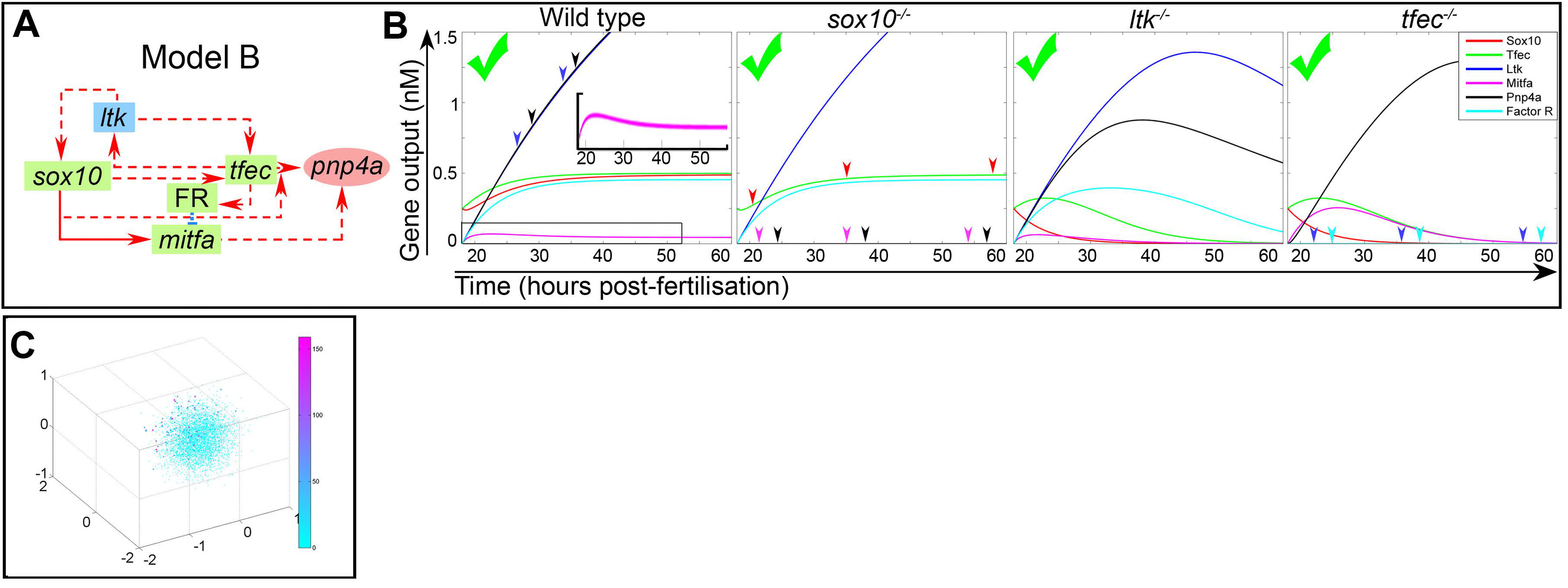
Detection of developing iridophores using expression of *tfec*. (A) Chromogenic WISH identifies *tfec* transcription in positions matching Iph positions at 72 hpf. (B) *tfec* is expressed in *sox10*- positive NCCs (region in brackets) at 18 hpf, which have downregulated *foxd3* but have not yet detectably activated early pigment lineage markers such as *ltk*, *mitfa* and *pnp4a*. (C) RNAscope reveals co-expression of *tfec* and *ltk* (arrows) in the ARPT during the timecourse of iridoblast specification, both in medially migrating cells at 24 hpf, and in dorsally located Ib(sp) and Iph at 30 hpf and 48 hpf, respectively. Using WISH at 24 hpf (D,E) *tfec* transcription is detectable in most or all cells of the multipotent premigratory NC domain (D, red line) and in a subset of cells of the posterior trunk located dorso-laterally to the spinal cord (D, vertical arrow) and in the medial pathway, between the somites and the notochord (D, horizontal arrow; Ei, arrows). Expression is only more weakly detectable in cells on the lateral pathway along the ARPT, between epidermal keratinocytes and somites (Eii, arrow). At 30 hpf (F,G) *tfec* is expressed in premigratory NCCs of the tail (red line) and in expected iridoblast positions, specifically dorsally located and medially migrating cells of the posterior trunk, the developing lateral patches and the eye (arrows). At 48 hpf (H) *tfec* is expressed along the dorsal, ventral and yolk sac stripes (vertical arrows), as well as in the lateral patches (arrowhead) and overlying the eye (horizontal arrow), in a pattern distinctive of differentiated iridophores. (I) RNAscope indicates presence of *mitfa* transcript in *tfec*-positive cells (arrows) migrating along the medial pathway in the ARPT at 24 hpf, but not in those located dorsally at 30 hpf, or at 48 hpf. In all three stages, *mitfa*+;*tfec*- cells are detectable (asterisks). All panels show lateral views, except dorsal views in (A; D,F insets). Head towards the left. RNAscope panels: single focal planes shown. e, epidermis; ICM, intermediate cell mass; LPs, lateral patches; no, notochord; RL, reflected light. Scale bars: (A,B,D,E,F,G,H) 50 μm; (C,I) 20 μm.

At 18 hours post fertilisation (hpf), premigratory NCCs reside along the dorsal trunk and tail (Fig 1B). Towards the posterior tail, these precursors are characterised by WISH as expressing markers such as *sox9b, sox10, snai1b* and *foxd3* (Dutton et al. 2001; Lister et al. 2006; Lopes et al. 2008) and likely correspond to fully multipotent early NCCs (eNCCs). At the same stage, more anteriorly (i.e. posterior trunk and rest of tail), *sox9b, snai1b* and *foxd3* are downregulated, while *sox10* is retained. Interestingly, *tfec* is expressed in premigratory NC cells throughout the trunk and anterior tail (Fig 1B), in a manner similar to *sox10*, even though fate-mapping of premigratory NCCs shows that only a relatively small subset of NCCs will generate iridoblasts (Dutton et al. 2001). At this stage neither the melanoblast marker *mitfa* nor two other early iridoblast markers *ltk* and *pnp4a* were detectable by WISH in *tfec*-positive NCCs of the trunk (Fig 1B) (Lister et al. 2006; Lopes et al. 2008). However, *mitfa* and *ltk* are activated widely by 22 hpf (Lister et al. 1999; Lopes et al. 2008), with *pnp4a* following soon after (Curran et al. 2010). We consider that these premigratory cells of the Anterior Region of the Posterior Trunk (ARPT; Fig 1B) which express *tfec*, but no longer *foxd3*, and which only later detectably upregulate other pigment markers, are multipotent iridophore progenitors corresponding to the proposed partially restricted pigment cell progenitor (Lopes et al. 2008). Given the spatiotemporal gradient of development that is so pronounced during the stages of NC development, throughout this paper we will largely focus on a readily-defined anatomical zone, the ARPT, when considering expression patterns at different stages, thus minimising the heterogeneity of the examined population of cells. The ARPT lies above the anterior yolk sac extension (YSE) (approximately the region of somites 9-11; bracketed in Fig 1B).

At 24 hpf, NCCs of the trunk have entered the medial and lateral migratory pathways, whereas less developed NCCs of the tail remain in premigratory positions. Although chromogenic WISH revealed maintenance of strong *tfec* expression in the premigratory NC domain, the majority of trunk NCCs located dorsal to the neural tube in the ARPT downregulate *tfec*, presumably due to cells becoming specified towards alternative lineages (Fig 1D). In this region, prominent *tfec* expression was retained in a small subset of precursors scattered over the spinal cord (Fig 1D), as well as in presumed iridoblasts migrating through the medial pathway (Fig 1Ei). *tfec* was only weakly detectable in cells entering the lateral migratory pathway, consistent with the medial migration pathway bias for iridoblasts noted previously (Kelsh et al. 2009) (Fig 1Eii). To further characterise the *tfec*-positive medially migrating cells, we used multiplexed fluorescent RNAscope to determine co-expression of *tfec* with the melanocyte lineage marker, *mitfa*. Importantly, *tfec-*expressing cells on the medial migration pathway were found to co-express *mitfa* (Fig 1I), leading us to distinguish these as iridophore progenitors which retain at least bipotency. In the context of this paper, we will designate such *tfec-*expressing cells on the medial pathway as specified iridoblasts (but equally they could also be considered specified melanoblasts, using the definition of Raible and Eisen (1994)). Other cells on the medial pathway were positive for *mitfa*, but displayed very weak or lacked expression of *tfec* (Fig 1I). We interpret these as having lost, or being on the way towards losing, iridophore potential, and are likely definitive melanoblasts. We did not observe *tfec*+;*mitfa*-cells in the APRT at this stage.

By 30 hpf, *tfec* transcript is present in scattered non-melanised cells along the dorsal posterior trunk and in medially migrating cells along the posterior trunk and anterior tail. Bilaterally patterned *tfec*+ premigratory NCCs were only detectable in the posterior tail (Fig 1F). In the APRT, co-expression analyses via RNAscope revealed that expression of *tfec* and *mitfa* has now resolved to be non-overlapping (Fig 1I); we now distinguish these cells as definitive iridoblasts, whether in the dorsal position characteristic of differentiated iridophores at later stages, or migrating on the medial migration pathway. Finally, we assessed *tfec* expression at 48 hpf, a stage when in live embryos differentiating, light-reflecting iridophores are distinguishable, interspersed along the dorsal, ventral and yolk sac stripes, occupying the lateral patches and overlying the eye. *tfec* expression was detectable by chromogenic WISH in all of these positions (Fig 1H). Co-expression analyses using RNAscope showed discrete expression of *tfec* and *mitfa* (Fig 1I), confirming that *tfec* expression at this stage definitively marked differentiating iridophores.

In summary, in addition to the fully multipotent early NCC state (eNCC), these marker studies clearly distinguished four sequential phases of iridophore development in WT zebrafish embryos (Fig 7): 1) premigratory NCCs presenting with widespread expression of *tfec*, but which have downregulated eNCC markers (for example, *foxd3*), interpreted as broadly multipotent pigment cell progenitors (*chromatoblasts,* Cbl); 2) scattered cells strongly maintaining *tfec* expression dorsal to spinal cord and on migration pathways, but also expressing *mitfa* and so interpreted as at least bipotent iridoblast progenitors (*specified iridoblasts,* Ib(sp)); 3) scattered undifferentiated cells in iridophore positions, showing rounded morphology and prominent expression of iridophore markers, but not *mitfa* (*definitive iridoblasts,* Ib(df)); and 4) discrete iridophore marker-expressing (and reflective in live fish) cells in characteristic definitive iridophore pattern (*mature iridophores,* Iph). We note that Ib(sp) and Ib(df) can only be distinguished in double WISH, and so will not be strictly distinguishable in many experiments, although from the above discussion it can be seen that at 30 hpf most such cells in the ARPT will be Ib(df). This characterisation provides a framework for assessment of expression patterns of other genes and for the interpretation of expression patterns seen in mutant embryos.

### *sox10* expression is maintained throughout development of the iridophore lineage

Although well-known as a key factor in iridophore specification and a key marker of multipotent NCCs (Dutton et al. 2001; Kim et al. 2003; Lopes et al. 2008), *sox10* expression has yet to be characterised in the iridophore lineage. We used both WISH and RNAscope to investigate the transcriptional dynamics of *sox10* during iridophore development. We imaged iridophores of live embryos at 72 hpf using reflected light, and subsequently detected *sox10* transcript in individual fish using chromogenic WISH. *sox10* expression was readily detected in all iridophores (e.g. in each of the dorsal, ventral and yolk sac stripes; Fig 2A-D). We then employed RNAscope to assess whether *sox10* expression was maintained throughout all stages of iridophore specification, as opposed to its becoming re-activated in differentiated cells. We found that all cells expressing the iridophore lineage marker, *ltk*, consistently co-expressed *sox10* at each of 24 hpf, 30 hpf and 48 hpf (Fig 2E-S). Therefore, just like *tfec*, *sox10* expression in premigratory multipotent NCCs (Fig 1B) is maintained throughout fate restriction to Ib(df) and their subsequent differentiation as iridophores.

**Figure 2.**
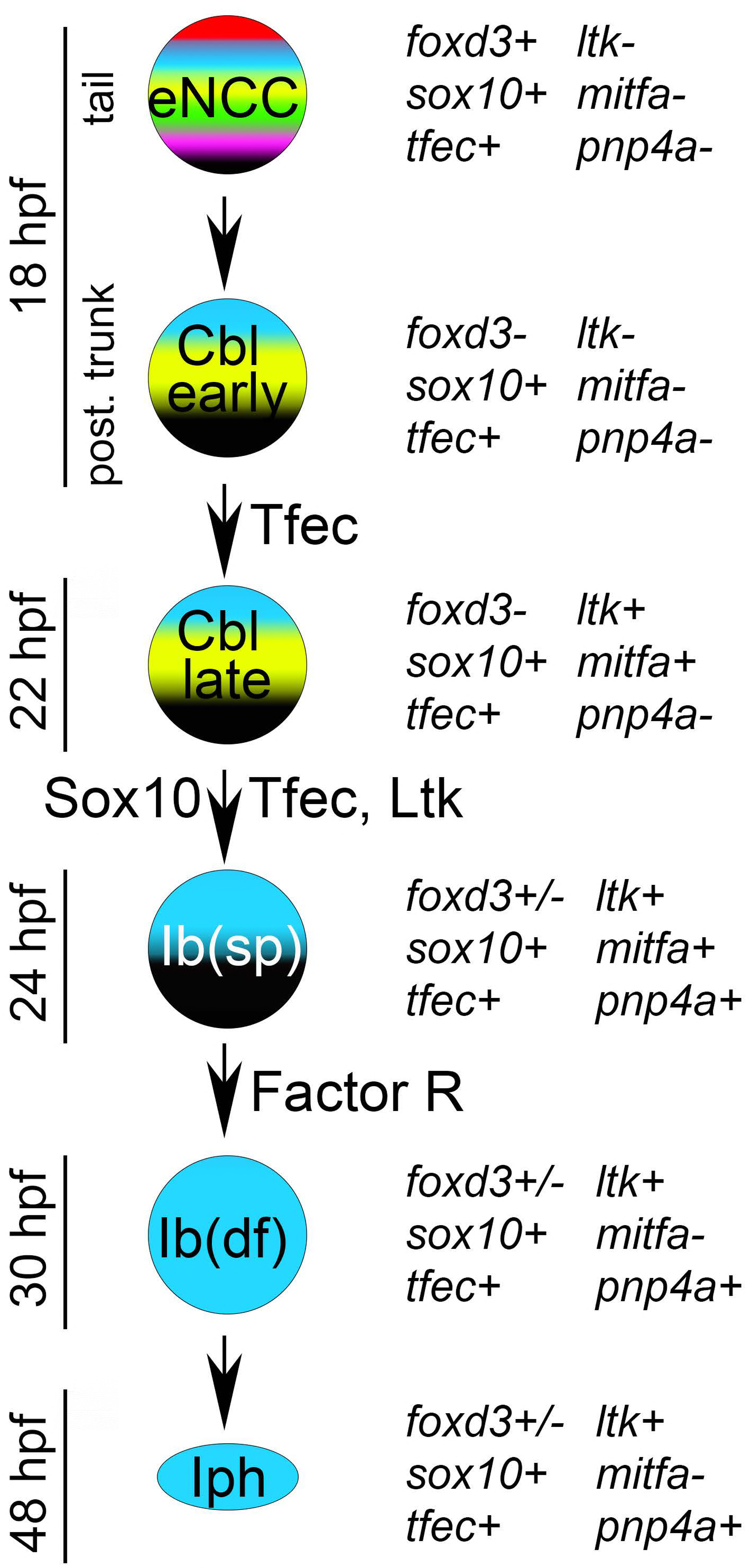
*sox10* expression is maintained throughout iridophore development. (A, B) lateral views of the anterior tail and (C, D) dorsal views of the trunk and anterior tail of individual embryos at 72 hpf, first imaged under reflected light (A, C) and then subjected to chromogenic WISH to detect *sox10* transcript (B, D). Differently coloured arrowheads point to individual iridophores expressing *sox10*. *sox10* is also detected in developing oligodendrocytes (B, arrow), Schwann cells (B) and in iridophores along the yolk sac stripe (A, B asterisk). *ltk*-positive cells detected via RNAscope (E, J, O arrowheads) all show *sox10* transcript (F, K, P; H, M, R arrowheads), at each of 24 (E-I), 30 (J-N) and 48 hpf (O-S). At 24 hpf, cells on the medial migration pathway are shown (I, boxed region). At 30 hpf and at 48 hpf, cells along the developing dorsal stripe are presented (N, S, boxed regions). (E-S): lateral views of single focal planes. (A-S): heads positioned towards the left. Sc, Schwann cells; pLLn, posterior lateral line nerve; no, notochord; RL, reflected light. Scale bars: (A-D) 100 μm; (E-H, J-M, O-R) 20 μm; (I, N, S) 50 μm.

### *sox10* maintains *tfec* expression in iridoblasts undergoing specification

Previous studies with *ltk* have concluded that iridophore specification fails and that pigment cell progenitors are trapped in a multipotent progenitor state (Cbl) in *sox10* mutants (Dutton et al. 2001; Lopes et al. 2008). We re-assessed this proposed role of *sox10* using both loss and gain of function assays. In *sox10* mutants, both at 24 hpf and at 30 hpf (Petratou et al., in prep; Fig 3A,B), *tfec* expression is prominently retained in the premigratory NCC domain, which unlike in WT siblings extends anteriorly throughout the embryo. These observations indicate that Sox10 function is not required for establishment of the *tfec-*positive multipotent progenitor. Importantly, *tfec* transcripts were undetectable in ventrally migrating iridophore progenitors, indicating a requirement for *sox10* to maintain *tfec* expression in a subset of cells (Ib(sp) and Ib(df)). By 48 hpf, we could not detect *tfec* expression in *sox10* mutant embryos (Fig 3E,F), consistent with apoptotic elimination of NC derivatives which fail to become specified, including progenitors for all chromatophore lineages (Dutton et al. 2001). To test the sufficiency of Sox10 for expression of *tfec,* we overexpressed WT Sox10 or a null mutant version by injection of mRNA into single cell stage WT embryos and assayed absolute *tfec* transcript levels using qRT-PCR at 6 hours post-injection. Our data clearly showed that functional Sox10, but not the null version of Sox10, ectopically activated expression of endogenous *tfec* (Fig 3O). Together these data show that Sox10 is not essential for initial activation of *tfec* in early NCCs, but it is required for maintenance of *tfec* expression as multipotent progenitors become specified towards an iridophore fate, i.e. for iridoblast fate specification.

**Figure 3.**
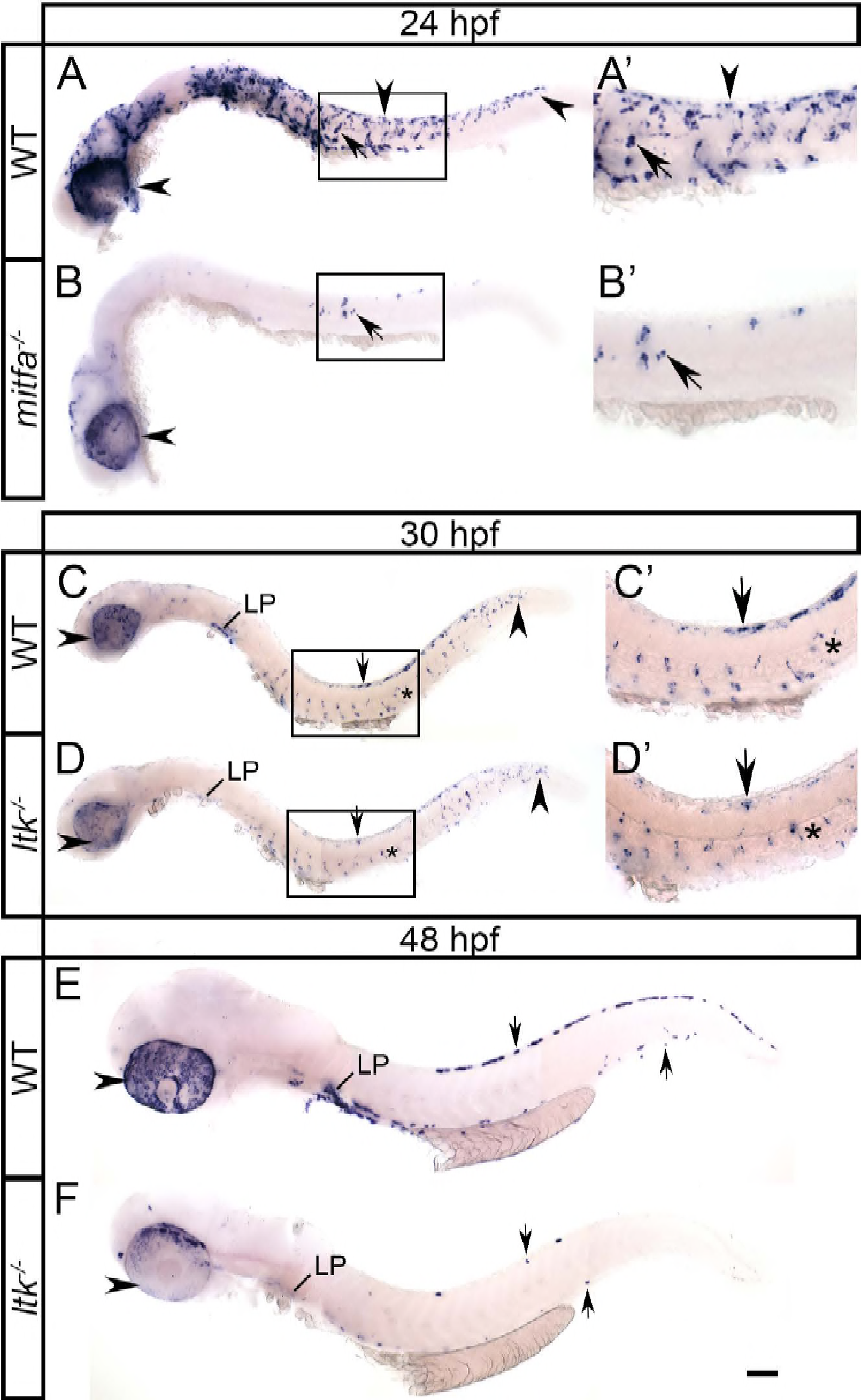
A *sox10*-dependent *tfec*/*ltk* positive feedback loop is required for iridophore specification. WISH to detect *tfec* (A-H) and *ltk* (I-L) expression at 30 hpf (A-D, I, J) and 48 hpf (E-H, K, L) in WT and mutant embryos, with quantitation (M,N). At 30 hpf, *sox10* mutants (B) lack *tfec* expression in medially migrating iridoblasts (A, arrows) and instead show a striking anteriorly expanded multipotent progenitor domain (B, arrowheads). Both *ltk* (C) and *tfec* mutants (D) display a reduced number of medially migrating Ib(sp) (horizontal arrows) at 30 hpf, while dorsally located Ib(sp) of the posterior trunk and tail (vertical arrows) are only significantly reduced in *ltk* mutants. At 48 hpf, *sox10* (F) and *ltk* (G) mutants lack *tfec*-labelling in the position of WT Iph (E, asterisks), while *tfec* mutants (H) display a reduced number of cells in these positions expressing *tfec* (H, asterisks; N). At both 30 hpf (I, J) and 48 hpf (K, L), *tfec* mutants lack *ltk* expression in Ib(df) (I, J, arrows) and in Iph (K, L, asterisks) locations, with the exception of rare escaper cells. (M) Quantitation of *tfec-*expressing iridophore lineage cells at 30 hpf. Counts for *tfec*-positive Ib(df) along the dorsal posterior trunk, the migration pathways, and the lateral patches are shown from left to right. (N) Quantitation of *tfec-*expressing cells along the posterior trunk and tail at 48 hpf. At this stage, scored cells are in Iph positions. *tfec* mutants display almost complete lack of *ltk*-positive cells both along the dorsal and the ventral stripes compared to their siblings, while *tfec* positive cells in both regions are partially reduced. (O) Overexpression of WT *sox10* mRNA results in an increased number of *tfec* transcripts, compared to overexpression of null *sox10* mRNA. A-L) Lateral views, head positioned towards the left. Scale bars: 100 μm.

### Tfec and Ltk generate a positive feedback loop

We next asked what roles Tfec and Ltk played in the iridophore GRN. Importantly, at both 18 and 24 hpf, we were unable to distinguish differences in *tfec* expression between *ltk* mutants and their WT siblings (S1 Table). Specifically, the premigratory NCC domain as well as specified iridoblasts in the posterior dorsal trunk and on the medial migration pathway of the trunk were unaffected in all examined embryos. Thus, *tfec* is activated in NCCs and is maintained at early stages of iridoblast specification, independently of Ltk activity. Nevertheless, from 30 hpf we observed a statistically significant decrease in the number of Ib(df) located in the dorsal posterior trunk of *ltk* mutants (Fig 3A,C,M), and by 48 hpf no cells expressing *tfec* were identifiable in the Ib(df) positions of the embryonic trunk in these mutants (Fig 3E,G).

Study of *ltk* expression in *tfec* mutants by chromogenic WISH suggests that Tfec function is crucial for *ltk* expression from the earliest stages onwards. At 24 hpf, approximately 25% of assessed embryos completely lacked *ltk* expression, with the exception of very rare escaper cells (Petratou et al., in prep.). This phenotype remained clearly identifiable at 30 hpf and persisted until at least 48 hpf (Fig 3I-L). Moreover, examination of *tfec* expression in *tfec* mutant embryos, readily distinguishable owing to lack of melanin pigment in the RPE (Petratou et al., in prep; Fig 3A,D insets), revealed a subtle but consistent reduction in the numbers of *tfec*-positive Ib(df) in mutants from 30 hpf (Fig 3A,D,M). Specifically, *tfec* mutants displayed a 35% and a 45% decrease in the number of *tfec-*expressing cells along the migratory pathways and in the developing lateral patches respectively, compared to WT siblings. Similarly, at 48 hpf, *tfec* mutant embryos (identified by the clear eye phenotype; Fig 3E,H insets) showed reductions in *tfec* expressing cells in the dorsal stripe and the ventral stripe to 58% and 45% of those in WT siblings (Fig 3E,H,N). Although the remaining *tfec-*expressing cells show a distribution consistent with their being Iph, we note that they lack both *ltk* expression and visible reflective platelets and hence cannot correspond to Ib(df), which are positive for *ltk* expression, nor to mature iridophores. We speculate that these cells represent an interesting state in which iridoblasts are trapped in a very early stage of their development, where *tfec* expression, but not other markers, continue to be maintained.

Taken together, our data strongly support the model that Tfec and Ltk function in a positive-feedback loop to maintain each other in specified iridoblasts, although *tfec* can be activated in this cell type independently of Ltk function.

### *pnp4a* is temporally regulated by variable activators

The gene *pnp4a* has been defined as an iridophore lineage marker, although it is expressed rather widely in NCCs and long before iridophores differentiate (Curran et al. 2010). RNA-seq analysis of gene expression in purified iridophores and melanocytes has shown that it is expressed at high levels not only in differentiated iridophores, but also, albeit at lower levels, in melanocytes (Higdon et al., 2013). We used chromogenic WISH studies to assess *pnp4a* expression in iridophore development and in various key mutants.

We first examined the WT expression pattern of *pnp4a*, and compared it to that of the iridophore and melanocyte lineage markers *tfec* and *mitfa*, respectively, at 24 hpf, 30 hpf and 48 hpf (Fig 4A-I). At 24 hpf, it is notable that the expression pattern of *pnp4a* strikingly resembled that of *mitfa*, rather than that of *tfec* (Fig 4A,D,G). Specifically, we see clusters of cells just posterior to the otic vesicle and numerous cells on the medial migration pathway in the expression patterns of both *mitfa* and *pnp4a*, although both these regions are only sparsely positive for *tfec* (Fig 4A,D,G and insets). By 30 hpf, this pattern is still detectable, and indeed now *pnp4a* transcripts are detectable in differentiating melanocytes clustered behind the otic vesicle, as well as in melanised cells of the head, in a pattern similar to *mitfa,* but not *tfec,* expression (Fig 4B,E,H and insets). In addition, at this stage, the pattern of *tfec* and *pnp4a* expression in both the dorsal posterior trunk as well as overlying the RPE showed strong similarities; *mitfa* transcript is absent from the latter region (Fig 4B,E,H). By 48 hpf, the pattern of *pnp4a* strikingly resembled that of *tfec*, with both transcripts detected in Iph positions, consistent with previously reported data (Curran et al. 2010), whereas *mitfa* was expressed in melanised cells of the head and of the dorsal, lateral and ventral stripes (Fig 4C,F,I). Considered together, our data suggest that while at later stages *pnp4a* is a definitive marker of differentiated iridophores, initially it is expressed widely in specified and differentiating melanoblasts.

**Figure 4.**
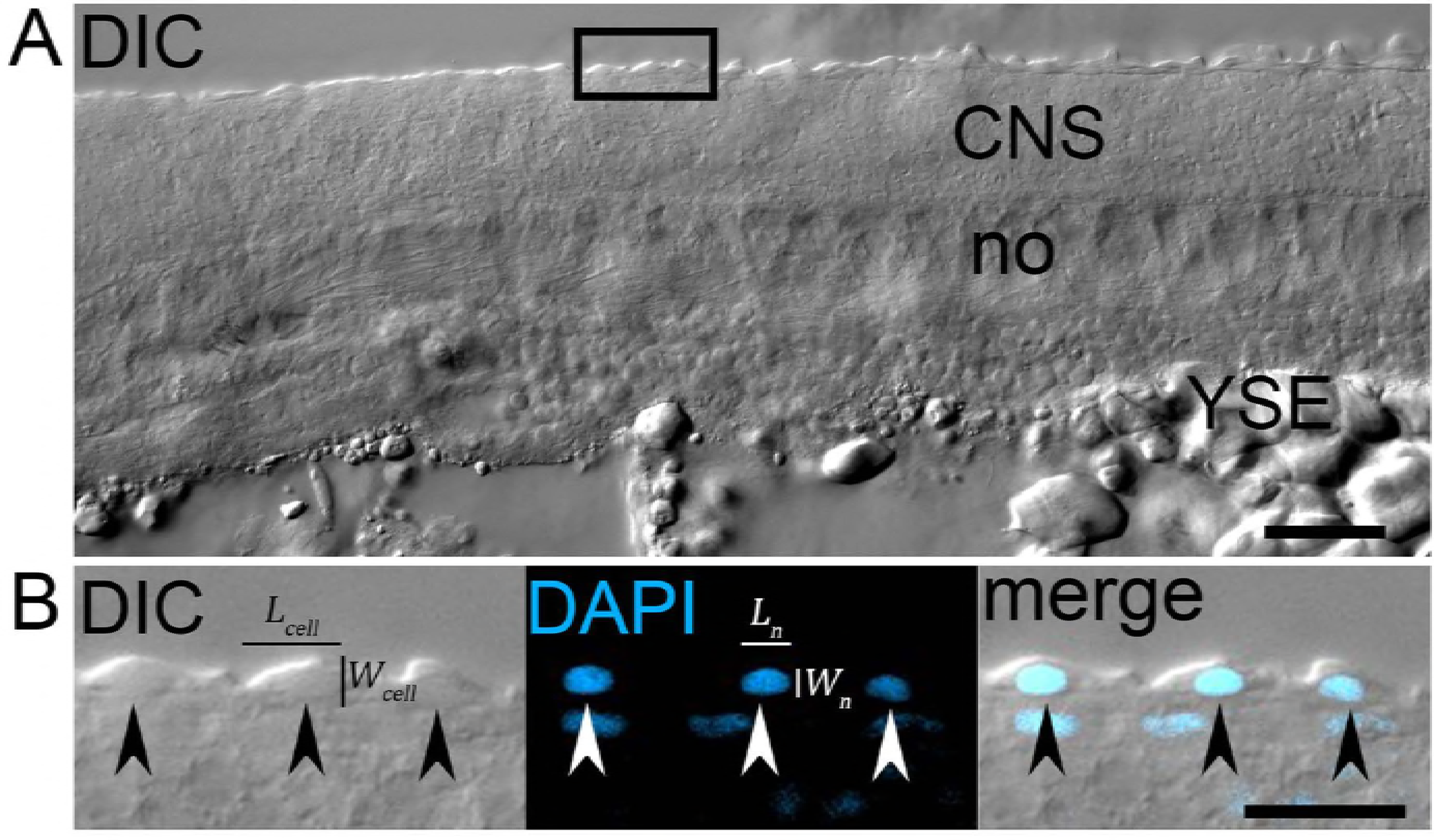
*pnp4a* is activated by *mitfa* in premigratory and migrating NC, and by cooperative action of *sox10* and *tfec* in iridoblasts. At 24 hpf and at 30 hpf, chromogenic WISH reveals strong similarities between the pattern of *pnp4a* and *mitfa* in migrating NC in the trunk and in cells posterior to the otic vesicle (A, B, D, E, black arrows), but which is distinct from *tfec*, which is present in more restricted groups of cells (G, H, blue arrows). At this stage, melanising cells in the head and anterior trunk show distinct expression of both *mitfa* and *pnp4a* (B,E, insets, black arrows). At 48 hpf, *mitfa* is expressed in melanised cells (F, black arrows), but *pnp4a* is not detectable in these melanocytes (C). From 30 hpf, some aspects of *pnp4a* expression are similar to those of *tfec* (B, H, blue arrows) and at 48 hpf both genes are expressed in Ib(df) locations (C, I, blue arrows). (J-Q) Mutant analysis. At 30 hpf, *mitfa* (M) and *tfec* (L) mutants retain only a subset of the WT *pnp4a* expression; remaining cells in the former display a Ib(df) pattern and in the latter a melanoblast pattern (M, L, arrows). *tfec* mutants lack *pnp4a* expression at 48 hpf, with the exception of rare escaper cells in iridophore positions (P, asterisks). Note that embryo in N, but not in O or P, was treated with PTU to inhibit melanisation; dark pigment in P is melanin. In *sox10* mutants, *pnp4a* is largely absent, although weak expression persists in a few premigratory NCCs (K, arrowheads) at 30 hpf, and in rare escaper cells in iridophore positions at 48 hpf (O, asterisks). (Q) qRT-PCR measurement of *pnp4a* expression after expression of Mitfa or Tfec in early zebrafish embryos. Overexpression of WT Mitfa results in ectopic activation of *pnp4a* in injected embryos at 6 hours post-injection, whereas mutant Mitfa (null) does not. Interestingly, neither WT nor mutant Tfec is sufficient to drive *pnp4a* expression at this stage. Fold activation is calculated following normalisation to *pnp4a* levels upon overexpression of GFP. *p*-values indicate the significance of mean fold change for each sample when compared to the mean of GFP, using a paired t-test. Lateral views, head towards the left. Scale bars: 100 μm. Inset scale bars: 50 μm.

This suggested that *pnp4a* expression might be regulated by both Tfec and Mitfa. We began by investigating *pnp4a* expression in *tfec* mutants and WT siblings. At 30 hpf, WTs showed prominent *pnp4a* expression along the dorsal and ventral posterior trunk and the migratory pathways across the trunk and tail (Fig 4J). In contrast, *tfec* mutants displayed partial loss of *pnp4a*-positive cells (Fig 4L). Specifically, compared to WT siblings, *tfec* mutants showed decreased numbers of cells (expressing relatively low levels of *pnp4a*) along the dorsal trunk, and have comparatively few cells both on the migration pathway and in the ventral trunk, mostly more anterior. This partial reduction was also observed at 24 hpf (Petratou et al., in prep.), and principally affected cells in the ventral trunk and premigratory NC. By 48 hpf, *pnp4a* expression in Iph locations was eliminated in *tfec* mutants (Fig 4N,P; Fig 3N). Thus, *pnp4a* expression in iridophores and in Ib(sp) is dependent upon Tfec, whereas the persistence of *pnp4a* expression in a subset of developing NC derivatives until 30 hpf suggested that *pnp4a* expression also depends on additional inputs.

Due to the striking similarity of their expression patterns, we investigated a possible interaction between *mitfa* and *pnp4a*. Interestingly, *pnp4a*-expressing cells were nearly eliminated in homozygous *mitfa* mutants, compared to their WT siblings at both 24 hpf (S1 Fig A-B’) and at 30 hpf (Fig 4J,M). Whereas in the former stage very few *pnp4a* expressing cells persisted along the posterior trunk of homozygous mutants (S1 Fig A-B’), in the latter a distinct group of dorsally located cells patterned in a Ib(df)-like manner along the posterior trunk and anterior tail region were retained. In contrast, medially migrating cells were almost absent (Fig 4M). These results indicated that *mitfa* is an important regulator of *pnp4a* in premigratory and migrating NC cells, but that *pnp4a* activation in Ib(df) is not affected. Thus, *pnp4a* expression appears to switch from Mitfa to Tfec-dependency during the transition from Cbl to Ib(df), and to be detectable transiently in all melanoblasts and early differentiating melanocytes.

We then asked whether Mitfa or Tfec were alone sufficient to drive *pnp4a* expression. We overexpressed each transcription factor, or a null mutant variant, in 1-cell stage WT embryos and measured *pnp4a* expression at 6 hours post-injection by qRT-PCR (Fig 4Q). As a negative control, we injected GFP RNA, allowing us to measure the relative (fold) change of expression between samples injected with GFP mRNA, compared to those injected with RNAs encoding WT or mutant. As expected, neither mutant Mitfa nor mutant Tfec altered *pnp4a* transcript levels, compared to overexpression of GFP. Interestingly, introducing WT Mitfa led to a statistically significant 4-fold increase (Fig 4Q). Surprisingly, however, overexpression of WT Tfec did not result in a statistically significant ectopic activation of *pnp4a* (Fig 4Q), suggesting that Mitfa, but not Tfec, is insufficient in this ectopic context to upregulate *pnp4a*.

Loss of function studies using *sox10* mutant embryos revealed that *pnp4a* was completely absent (Fig 4K), and that the gene remained inactive throughout the investigated developmental time-course (Fig 4O). We concluded that *sox10* function was directly or indirectly required for all aspects of *pnp4a* expression, including the *tfec*-dependent *pnp4a* upregulation to occur. More broadly, we conclude that *pnp4a* regulation is more complex than previously assumed and that it should not be considered a definitive marker of the iridophore lineage at early stages.

### A preliminary GRN governing iridophore development

Bringing the above interactions together, we propose a preliminary iridophore GRN, comprising model A (Fig 5A-C). We use solid lines to describe known direct interactions and dashed lines to indicate interactions where their nature is unknown. Sox10 has been previously shown to bind directly to the promoter of *mitfa* in zebrafish, and to activate its expression (Elworthy et al. 2003), we thus include that interaction. Furthermore, Ltk relies on intracellular cascades and effector transcription factors to activate gene expression, thus its input is always indirect. However, for the remainder of the interactions it remains unclear whether Sox10, Tfec or Mitfa bind directly to the promoters of downstream genes.

**Figure 5.**
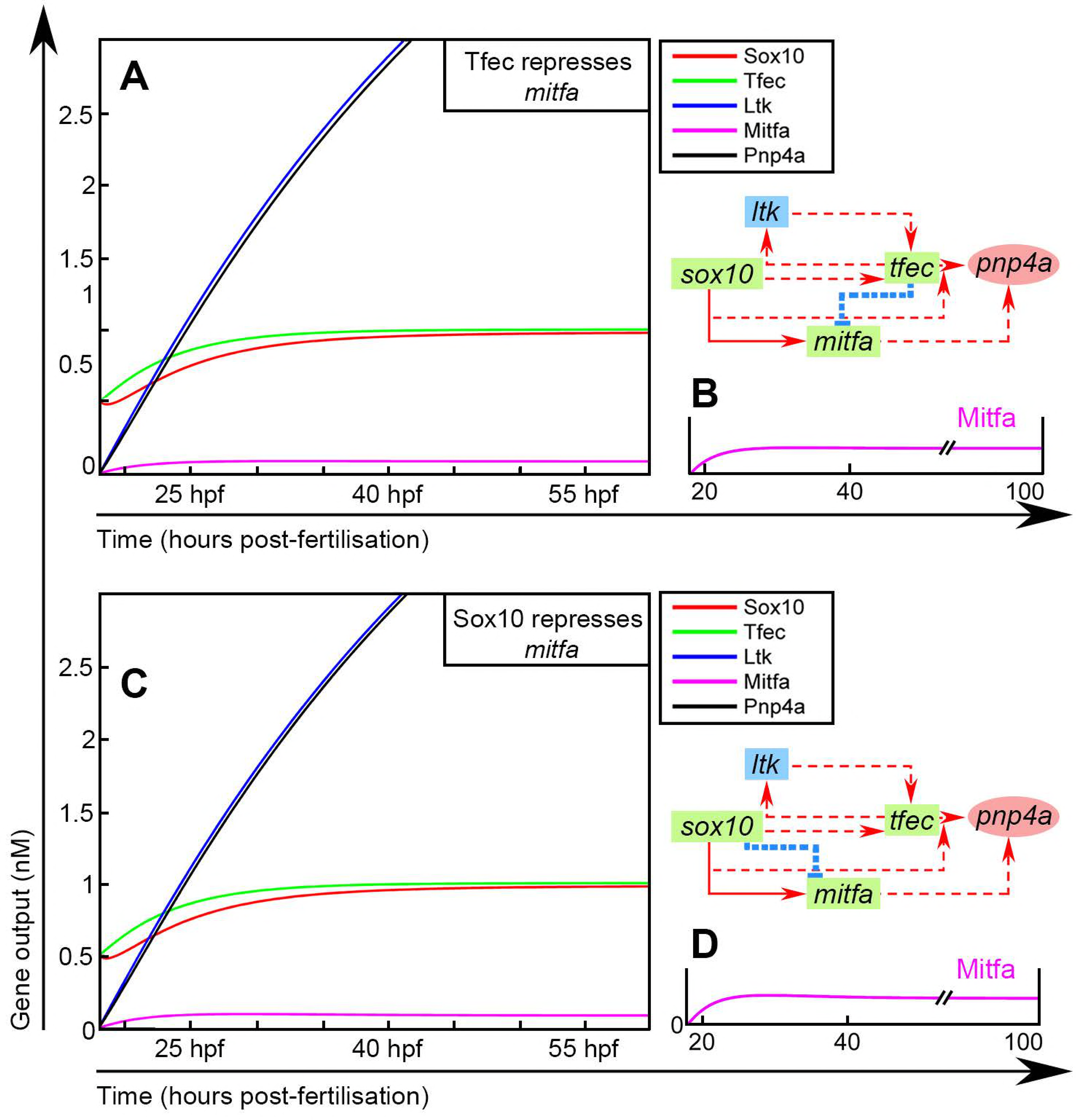
Mathematical modelling and refinement of the preliminary iridophore GRN. Graphical representation, simulation outputs and PCA analyses of Monte Carlo results for models A1 (A,D,G), A2 (B,E,H) and A3 (C,F,I)). In the schematics (A-C), green rectangular nodes are used for transcription factor-coding genes, blue rectangles for transmembrane receptors and red circles for enzymes. Transcriptional activation is indicated by red arrows. Solid edges represent known direct interactions, whereas dashed lines indicate known indirect or interactions where their nature is unknown. These diagrams are mathematically described using ODEs, the numerical solutions of which indicate how the gene expression dynamics progress in WT and mutant scenarios (D-F). In the simulation outputs (D-F), coloured lines represent the change of concentration (nM) for each gene output during the course of iridophore development (hpf). The simulation considers the developing iridophore lineage starting from *tfec+;sox10+;ltk-;mitfa-;pnp4a-* NCCs of the ARPT at 18 hpf. Qualitative assessment of simulation outputs is indicated as follows for each gene. Green tick: modelling predictions match experimental observations. Crosses: prediction for a specific gene (indicated by colour of the cross, matching gene reference colour as indicated in the legend) is refuted by experimental evidence. (G-I) PCA plots for models A1-A3. All trials using model A1 fail the set criteria, thus scoring between 0-0.2 (G, turquoise spots). Under model A2, a small proportion of parameter sets scores within the range of 25-55 (H, dark blue and purple spots) and very few trials score between 55 and 75 (H, magenda spots). Under model A3, performance improves notably, with more trials scoring between 25 and 55 (I, dark blue and purple spots), and others scoring between 55-80 (I, magenda spots)

We propose three distinct variants of Model A, distinguished by the mechanism of *sox10* maintenance in iridoblasts and iridophores (Models A1, A2 and A3 (Fig 5A-C)). In A1, *sox10* maintenance occurs through an autoregulatory positive feedback loop (Fig 5A), which maintains *sox10* expression from premigratory progenitors (Fig 1B). In A2, the presumed iridophore master regulator, *tfec*, is directly or indirectly responsible for *sox10* activation in the context of iridoblasts (Fig 5B). Finally, model A3 proposes that *sox10* maintenance is dependent upon Ltk signalling, independent of Ltk’s action in maintenance of Tfec expression (Fig 5C).

### Mathematical exploration of the preliminary iridophore GRN

These three Model A variants all share positive feedback loops between Sox10, Tfec and Ltk, and whilst biologically distinct, it is not obvious intuitively how they could be distinguished without detailed investigation of transcriptional regulatory mechanisms. However, like others, we have previously demonstrated the value of simple predictive mathematical modelling of GRNs in developing a robust understanding of their biological implications (Greenhill et al. 2011). Thus, we utilised mathematical modelling of each model A variant, to assess more rigorously whether they could be distinguished. If so, we wished to identify the model offering highest predictive power, i.e. the network that was best able to recapitulate the experimentally observed gene expression dynamics in the iridophore lineage.

We generated systems of ordinary differential equations (ODEs) describing the interactions in each of the proposed networks (see S1 text). The changes in the expression of each gene over time were determined using an ODE which incorporated all activatory and repressive influences from other members of the network, as well as a term for degradation of the gene’s protein product. The model aimed to capture the average output (nominally as protein product, assuming direct correlation with transcript production) of each gene in a homogeneous group of cells at a given time. The necessary parameter values characterizing the regulatory dynamics (mRNA maximum production rates (g), protein degradation rates (d), dissociation constants for transcription factors binding (K)) were chosen following exploration of existing literature to identify physiologically relevant values (see S1 text; S2 Table).

We solved the systems of ODEs numerically in MatLab. To define initial conditions (here at t = 18 hours), we chose the population of premigratory NCCs occupying the dorsal ARPT at 18 hpf. Using chromogenic WISH, *sox10* and *tfec*, but not *ltk*, *mitfa* or *pnp4a* transcripts were detectable in this population of cells (Fig 1B), allowing us to approximate the initial conditions for our simulations. As time proceeds, the simulations were tested for their ability to broadly replicate the changes in gene expression of *tfec*- expressing cells as they transition through the stages of Ib(sp), Ib(df) and then Iph. *In vivo*, differentiating iridophores are observable from 42 hpf and prominent by 48 hpf. To account for inaccuracies in our default parameter sets (see S1 Text; S2 Table), we allowed computational simulations to progress until 60 hpf, thus helping to ensure that any biologically meaningful steady state could be successfully reached. Specifically, in the WT context, it was crucial that Sox10 and Tfec stay upregulated in mature pigment cells, i.e. reach a positive steady-state. Similarly, Ltk and Pnp4a concentrations should increase and reach a positive steady-state. Mitfa levels should initially rise rapidly, reflecting the widespread expression of *mitfa* in all pigment cell progenitors (Curran, Raible, and Lister 2009), but should then drop to a distinctly lower level at differentiation stages (this work). We note that the Mitfa concentration is not required to attain zero, but simply to drop to a lower steady-state value; given the expectation that *in situ* hybridisation techniques have a ‘detection threshold’, we consider that this final lower value would reflect expression levels undetectable by our detection methods in differentiated iridophores, although they would still be measurable by microarray in pooled isolated iridophores (Higdon, Mitra, and Johnson 2013). As a further test of each of Models A1, A2 and A3, we used MatLab to predict gene expression changes in the context of different mutant scenarios, when function of Sox10, Ltk or Tfec were individually ablated *in silico* (Fig 5). In the *sox10* mutant context we asked that Sox10, Tfec and Ltk acquire positive values, as expression has been identified in trapped chromatoblasts (Dutton et al. 2001; Lopes et al. 2008, this work), however at no point do either Mitfa or Pnp4a become upregulated. Loss of Ltk function was required to predict initial rise of Ltk, Tfec and Pnp4a concentrations (at approximately 24-30 hpf), followed by gradual downregulation to undetectable levels. Similarly, loss of Tfec function should be accompanied by a peak and subsequent decline of Tfec, Sox10 and Pnp4a concentrations within 30-50 hpf. Ltk should never become detectable in the developing iridophore population in this context.

These simulations showed that regardless of the interaction underlying *sox10* maintenance, in the WT context all iridophore markers were appropriately upregulated in the course of iridophore development, consistent with biological observations. In all three models, expression of the melanocyte marker, *mitfa*, a direct target of Sox10 (Elworthy et al. 2003), was predicted to be upregulated and then maintained in the iridophore lineage (Fig 5D-F). These predictions are in contrast to previously published experimental data, showing that maintenance of *mitfa* is restricted to melanocytes (Elworthy et al. 2003; Greenhill et al. 2011), although lower level *mitfa* expression has been detected by RNA-seq in differentiated iridophores (Higdon, Mitra, and Johnson 2013). We used RNAscope to assess directly the relative changes in *mitfa* expression in the iridophore lineage. Even with this technique, notably more sensitive than conventional chromogenic WISH, we confirmed that co-expression of *mitfa* with the iridoblast marker, *tfec*, occurred at 24 hpf, but such overlap was not detectable at 30 hpf and at 48 hpf (Fig 1I). Thus, all three models failed to correctly predict the expected initial peak, followed by downregulation, of *mitfa* expression in the iridophore lineage. These observations are readily explained by the absence of a mechanism for repression of melanocyte fate in our model; we explore this later.

For all three versions of model A, simulation of loss of Sox10 function appropriately predicted maintenance of *tfec* (Fig 3B) and of *ltk* (Lopes et al. 2008), consistent with observations that *tfec+;ltk+* progenitors remain trapped in the dorsal trunk and tail. Likewise, they appropriately predict the failure to upregulate both *mitfa* (Elworthy et al. 2003) and *pnp4a* (Fig 4K). However, it has been previously shown that dorsally trapped progenitors continue to express *sox10* upon loss of Sox10 function (Dutton et al. 2001), a feature predicted successfully by models A2 and A3, but not by A1. Similarly, computational implementation of *tfec* loss of function revealed that model A1 did not generate biologically accurate predictions, whereas models A2 and A3 performed better. Specifically, Model A1 with simulated loss of Tfec function did not result in the experimentally observed lack of *ltk* and gradual downregulation of both *tfec* and *pnp4a* expression (Fig 3D; Fig 4L). In models A2 and A3, *ltk* expression was correctly predicted to remain undetectable throughout iridophore specification and differentiation, while *tfec* and *pnp4a* were gradually diminished. Finally, *in silico* inhibition of Ltk signalling in models A1 and A2 failed to predict the experimentally observed initial activation, followed by downregulation, of *ltk* (Lopes et al. 2008), *tfec* (Fig 3C) and *pnp4a* (S1 Fig C-F) expression as iridoblasts differentiate into iridophores. Model A3, however, successfully predicted gradual elimination of iridophore marker gene expression in the lineage, more accurately reflecting the current experimental observations.

Based on the above observations, we conclude that Model A3 has the highest degree of predictive power using the default parameter set. These parameters were chosen based on ranges indicated from the literature as physiologically relevant, nevertheless the exact values were assigned somewhat arbitrarily. We, therefore, conducted an unbiased assessment of whether the experimentally set output requirements, as outlined above, could be achieved using alternative sets of parameter values in any of the models. To that effect, we designed a Monte Carlo algorithm able to randomly assign parameters drawn from a pre-assigned range, spanning two orders of magnitude from 5x higher than the physiological mean value to 5x lower than that value. For each model, the outputs for each of 20,000 combinations of parameters were scored computationally by a suitably designed scoring function, according to our set of qualitative criteria (see S2 Text), and which took into account that only qualitative expectations of gene regulatory dynamics could be tested. Principal component analysis (PCA) was used to visualise the scoring results within parameter space and to compare the three models’ respective capacities to reproduce experimental observations (Fig 5G-I). Interestingly, all outputs derived from randomly assigning parameter values in the set of equations representing model A1 failed to predict crucial aspects of the biology, thus consistently achieving zero scores (Fig 5G). Model A2 was found able to predict those features correctly, although only limited subsets of parameters achieved admissible outputs (Fig 5H). Model A3 performed similarly to Model A2, except that it generated predictions broadly consistent with the known biology for a wider range of parameter combinations (Fig 5I).

### Repression of *mitfa* in the iridophore lineage requires an unknown Factor R

Although the analysis of the model A variants identified a more favourable model (A3) for most aspects, none of the model A alternatives were able to reproduce the expected Mitfa dynamics (i.e. sufficient downregulation of Mitfa in differentiating iridophores), while simultaneously maintaining relatively high outputs of iridogenic gene products (Fig 5; S5 Fig A-C). We used model A3 as a starting point to improve this aspect of the iridophore GRN. Since repression of *mitfa* in the iridophore lineage has not thus far been investigated, we asked which interactions would be able to produce appropriate outputs when mathematically implemented. After testing predictions of alternative models with our default parameter set (S3 Fig), we concluded that upregulation in the iridophore lineage of an unknown *mitfa* repressor, which we termed factor R, was crucial. The resulting Model B (Fig 6A) incorporated Tfec-dependent activation of factor R, which our implementations suggested should be absent in the multipotent progenitors of the ARPT at t = 18 hours. Manually adjusting the parameters in the system of ODEs describing Model B revealed that the experimentally determined rise and drop of *mitfa* expression in our group of cells could be achieved using the default parameter set (Fig 6B), and even enriched with alternative parameter values, within the determined physiologically relevant range (S5 Fig E,G). Random assignment of parameters and algorithmic scoring of respective outputs (see S1 Text) indicated that, of all the tested models, model B best reflected experimental observations regarding gene expression dynamics. Specifically, compared to models A1-A3, a broader range of model B trials achieved high scores, with absolute values higher than those attainable through models A1-A3 (Fig 5; Fig 6). Notably, model B (derived from model A3) consistently scored higher than a designated alternative model B(2), which was derived by introducing factor R into model A2 (S5 Fig F-G).

**Figure 6.**
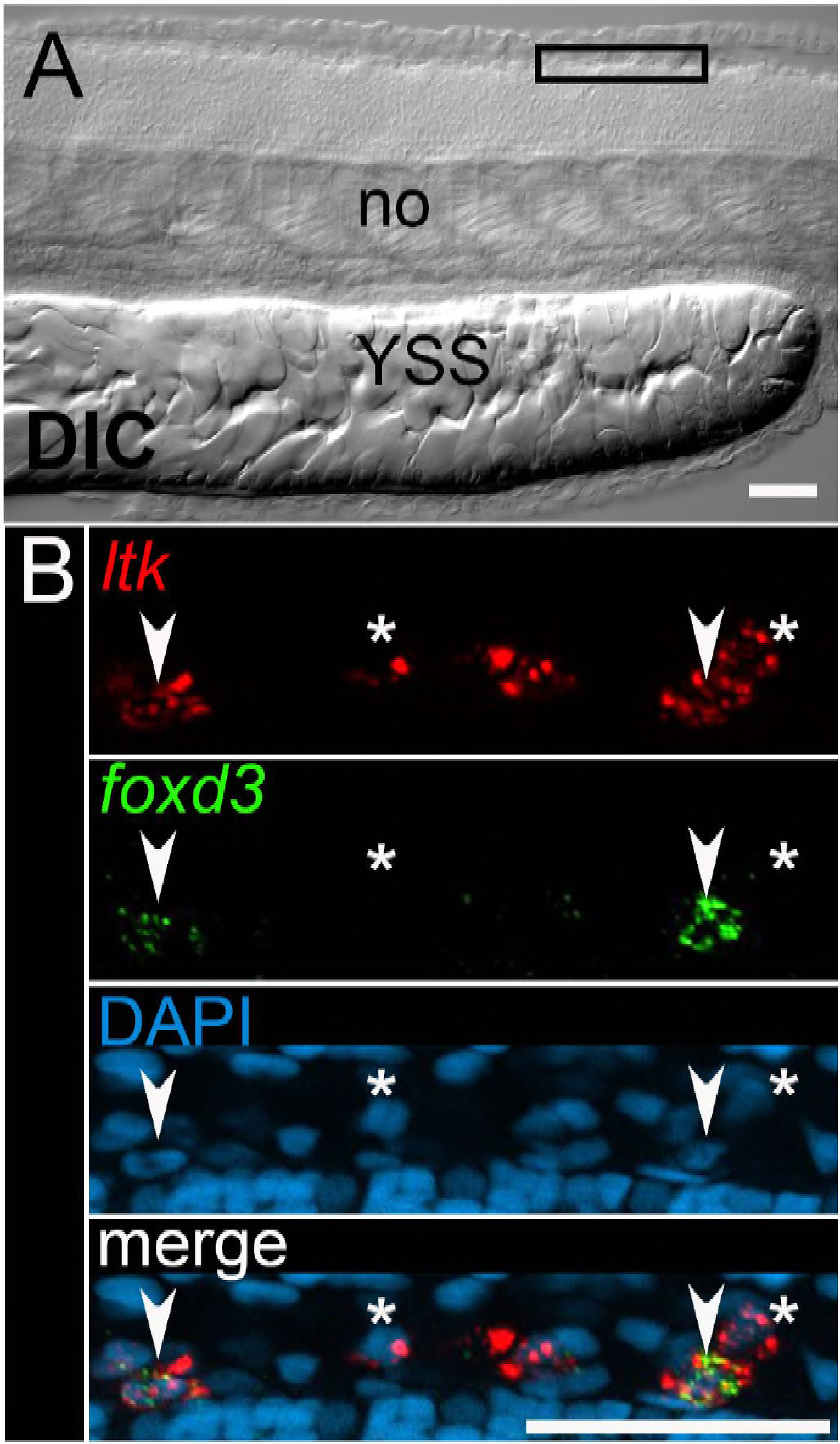
Model B accurately reflects observed gene expression dynamics. (A) Graphical representation of model B, where blue (blunt-ended) edges indicate transcriptional repression. (B) Simulation outputs. (C) PCA plot for model B. A large proportion of random parameter trials yields scores between 50-100 (dark blue and purple spots), while several trials achieve scores from 100 to as high as 150 (purple/magenda spots).

We considered Foxd3 as a candidate for factor R, as the transcriptional regulator has been previously implicated in *mitfa* repression (Curran et al. 2010). Our modelling predicted that, in the WT context, factor R should be expressed in undetectable levels in the ARPT at 18 hpf, and should then be upregulated in all developing iridophores, before reaching a stable plateau at differentiation. We tested these predictions for *foxd3* using RNAscope. Surprisingly, we detected only low levels of *foxd3* transcript, and these in only 50% of *ltk*-positive iridophore lineage cells at each of 24 hpf, 30 hpf, 36 hpf and 48 hpf (S4 Fig). Hence, we conclude that *foxd3* is unlikely to function as factor R in our modelling of iridophore development.

## Discussion

In previous work we used an iterative process of experimental genetics and mathematical modelling to develop a robust core GRN for the zebrafish melanocyte. Here, we extend that approach to establish a core GRN for the iridophore, a second pigment cell-type that shows a close developmental genetic relationship with the melanocyte, and which has been proposed to derive from a shared bipotent progenitor (Curran et al. 2010).

In the course of our experimental analysis, it soon became clear that iridophore-related genes often showed multiphasic expression, being detectable in differentiated iridophores, but also in much earlier, even premigratory stages of NC development. We had first noted this in our study of *ltk* expression (Lopes et al. 2008), but here we showed that *tfec, pnp4a* and *sox10* behave similarly. This same phenomenon has been documented, but not emphasised, in the case of melanocyte development, with *mitfa* being expressed initially in almost all NC cells (Curran, Raible, and Lister 2009), but it is less clear whether other melanocyte-specific genes present with similar biphasic expression. Our use here of the RNAscope assay, readily allowing highly sensitive detection and quantitation of co-expression, reveals that early markers of fate specification of different cell-types (e.g. *mitfa* and *tfec*) may be initially co-expressed. This reflects the distinction between fate specification, when a cell is beginning to show characteristics of a specific lineage, and commitment, when it has stably adopted that fate at the expense of alternative ones. These considerations, plus the standard limitation that we usually examine only one or two markers at once, resulted in us attempting to standardise our assessment of gene expression patterns, taking account of not only marker expression and levels of expression, but also cell location and cell morphology. This led to an explicit working model of stages in iridophore development from early NCCs (Fig 1 and Fig 7). This model is broadly consistent with the current progressive fate restriction model of NC development. However, the experimental restrictions noted above mean that we can, at best, assess minimal levels of potency: where we see overlap of expression of key genes for different fates, we interpret this as reflecting the cell having potential for *at least* these fates. Analysis of *ltk* expression in *sox10* mutants (Lopes et al. 2008, Nikaido et al., in prep.), interpreted in the light of our detailed studies of the mutant phenotype, including single cell fate-mapping of NC (Dutton et al. 2001; Subkhankulova et al., in prep.), led us to propose that premigratory NCCs in the trunk and tail go through a multipotent pigment cell progenitor phase, that we refer to here as the Cbl phase (Fig 7). Combining the RNAscope and WISH data presented here with those studies and our similar observations for other markers, provides support for this interpretation, since premigratory cells expressing *tfec* or *ltk* do not express markers of fully multipotent eNCCs (*e.g. foxd3,* but also *snai1b* and *sox9b*). We further distinguish two phases to this Cbl stage, with cells initially expressing *tfec*, but not *ltk, mitfa* nor *pnp4a* (which we designate early Cbl cells), before rapidly turning on all these genes (becoming late Cbl cells; Fig 7).

**Figure 7.**
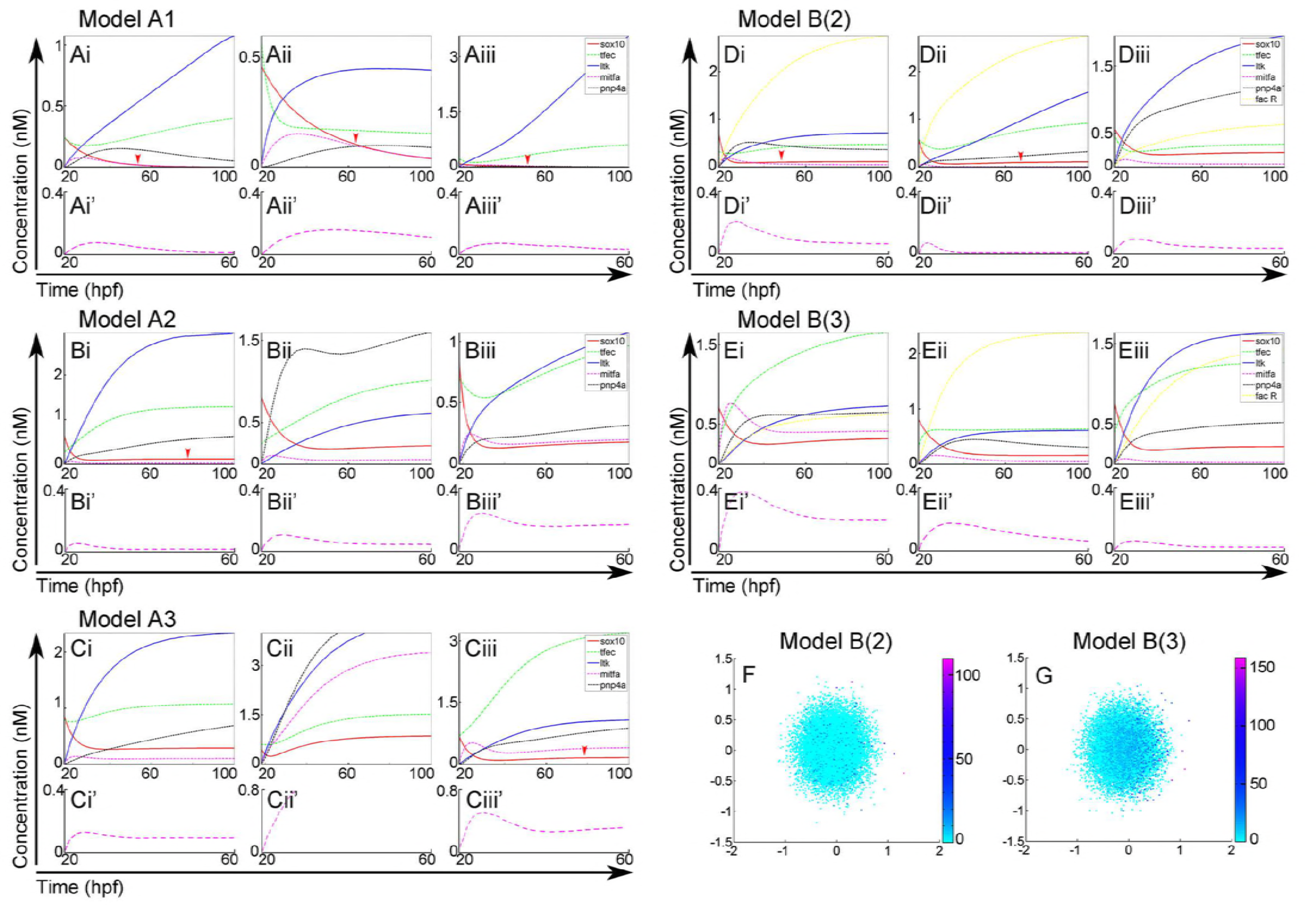
Progressive fate restriction model for iridophore development from eNCCs. We interpret our gene expression and mutant data in the context of a progressive fate restriction model, identifying a series of intermediate stages: early and late Cbl, Ib(sp), Ib(df) and Iph. For each cell type, the relevant stage and trunk region is stated on the left and the characteristic genetic signature on the right. eNCC, early NCC; Cbl, chromatoblast; Ib(sp), specified iridoblast; Ib(df), definitive iridoblast; Iph, iridophore.

A crucial step is then the establishment of a positive feedback loop between Ltk and Tfec, which drives the maintenance of iridophore specification state. Our findings build on the established role of Ltk signalling, linking it to a key transcription factor for iridophore fate specification, Tfec. Tfec is a close homologue of Mitfa, so it is intriguing that it seems to have a similarly central role in iridophore fate specification as does Mitfa in melanocyte development. However, Tfec is expressed much earlier than *mitfa* and *ltk* in NC development, being detected widely in early NCCs (Lister et al. 2011; Petratou et al., in prep.). Although the upstream regulation of *tfec* in early NCCs is only starting to be elucidated (Petratou et al., in prep.), we here show that Ltk signalling is required to maintain Tfec in a *subset* of cells, setting them aside as Ib(sp). However, it is important to note that these cells are initially co-expressing *mitfa*, consistent with their being at least bipotent progenitors of both melanocytes and iridophores. Intriguingly, we show that one early function of Tfec is to activate *ltk* expression in premigratory NCCs, in what we consider to be a first step in chromatophore fate restriction (Fig 7), distinguishing the multipotent Cbl from the fully multipotent eNCC.

Importantly, this work highlights the ongoing role of Sox10 in iridophore development. In melanocytes, Sox10 acts together with Wnt signalling to establish *mitfa* expression, but *sox10* is then strongly downregulated, and indeed maintenance of expression is thought to promote multipotency and delay differentiation (Greenhill et al. 2011). In the iridophore lineage, loss of *sox10* function results in failure of iridophore fate specification, suggesting that there are strong parallels between the genetic mechanisms of melanocyte and iridophore fate specification. Surprisingly, in contrast to the regulatory dynamics taking place in the melanocyte lineage, our data indicate an ongoing role for *sox10* over the course of iridophore development, with expression of the gene being readily detectable by RNAscope in specifying iridoblasts and by WISH in mature iridophores. It will be interesting therefore to explore the molecular basis for repression of alternative fate choices that allows *sox10* expression (which is strongly associated with NCC multipotency;Paratore et al. 2002; Kim et al. 2003; Kléber et al. 2005; Kelsh 2006) and iridophore fate commitment to proceed hand-in-hand.

Our experimental data showed that *sox10* expression needed to be maintained, but the close regulatory relationship between the Ltk-Tfec feedback loop and Sox10 made it difficult *a priori* to distinguish three variants: 1) sox10 autoregulation, or 2) input from the Ltk-Tfec loop through Tfec (or a downstream target of Tfec), or 3) through Ltk independent of Tfec. Intuitively, the impact of these three distinct modes is difficult to decipher, but an unexpected outcome of the mathematical modelling was the realisation that the behaviour of the GRN was quite different under these models. Our simulations, supported by unbiased random sampling via a Monte Carlo approach clearly suggested that the third model (Model A3) was superior to the alternatives, in that it most readily and robustly led to a predicted pattern of gene expression most closely mimicking that observed experimentally. This nicely illustrates the benefits of simple predictive mathematical modelling in rigorous assessment of GRNs.

Our observations also revealed an unexpected complexity to the regulation, and thus the likely role, of *pnp4a* in pigment cell development. Although the gene has been considered a definitive marker of the iridophore lineage (Curran et al. 2010), our study reveals complex regulation of *pnp4a* in premigratory and migrating NCCs, by both Tfec and Mitfa, as well as by Sox10. During these stages of fate specification and early differentiation *pnp4a* is best considered a marker of both specified melanoblasts and specified iridoblasts (which as we have shown likely include many shared cells), although we also confirm that at later stages by WISH at least it is a definitive marker of the iridophore lineage. This gene encodes purine nucleoside phosphorylase 4a, an enzyme converting guanosine mono-phosphate to guanine (Higdon, Mitra, and Johnson 2013). We note that in medaka the *guanineless/pnp4a* gene mutant phenotype is a pronounced reduction of iridophore reflectivity, consistent with its proposed enzymatic role in generating high concentrations of guanine in iridophores to allow reflecting platelet formation (Kimura, Takehana, and Naruse 2017). The gene’s role in melanoblasts remains unclear.

One significant innovation in our implementation of the mathematical modelling in this study over its use in the Greenhill study, is in our assessment of parameter values. One well-known problem with mathematical modelling is that as GRNs increase in complexity the number of parameters increases rapidly. In most *in vivo* systems, the absolute values of these parameters cannot be easily measured, leading to considerable uncertainty about the resulting simulations and their validity. To overcome this drawback in our modelling here, we restricted the values of all parameters to physiologically relevant ranges, based upon published measurements, choosing values that we consider to be best estimates based upon the same or similar molecular interactions. Furthermore, we employed an unbiased Monte Carlo sampling to ask explicitly how significant changes to those parameters might be for the output of the models. For example, we asked whether randomly varying the originally assigned ‘sensible’ parameter values, implementation of which had resulted in a particular model’s predictions broadly matching experimental observations, might render the subsequent modelling outputs radically different and thus the model less convincing. Alternatively, we aimed to confirm that any model that gave inaccurate predictions with the originally assigned parameter set remained incapable of predicting the experimentally observed gene regulatory dynamics even with different parameter sets. We emphasize that the method presented here is particularly suitable to all modelling attempts where experimental data is qualitative in nature and limited to few developmental time-points due to technical restrictions. In this sense, it addresses an important drawback, typical in Systems Biology model reconstructions, when both the topology of the network and the parameters are unknown. In standard fitting procedures the topology is assumed, and the optimal parameter set is chosen so as to minimise the discrepancy between the theoretical predictions and the (quantitative) experimental data. This approach of course fails when a quantitative dataset is not available. The method described here aims to cover those situations where only a limited, non-quantitative set of data on dynamical behaviours is available, and attempts to assess different network topologies in a broad spectrum of parameter values. By augmenting the model GRN with the presented scoring functions, our Monte Carlo screening algorithm allowed us to rigorously explore the proposed model variants and to compare their predictive powers under a broad set of physiologically-relevant parameter values. This approach renders the process of either validating or refuting model variant considerably more objective.

The assessment of our GRN by mathematical modelling revealed a key feature, one that will be crucial when integrating the melanocyte and iridophore GRNs, namely the factor (factor R) repressing *mitfa*, and thus melanocyte fate, in the iridoblasts. A series of published experimental observations, including the *foxd3* mutant phenotype, with partial loss of iridophores, partially rescued in *foxd3;mitfa* double mutants (Curran et al. 2010), had led to the proposal that FoxD3 might have such a role. Our modelling indicates that factor R is required throughout iridophore development, including into differentiation phases. As a test of the suitability of *foxd3* for this role, we assessed expression in the iridophore lineage throughout a developmental time-course using both conventional WISH, as well as RNAscope. Previous analyses using a *foxd3*:*gfp* transgenic line have suggested that Foxd3 is expressed in mature iridophores (Curran, Raible, and Lister 2009). Here we used RNAscope to assess co-expression of *foxd3* with the iridophore lineage marker *ltk*. This technique is both highly sensitive and quantitative, thus much better suited for sensitive detection of co-expression, with the added advantage that rapid turnover of mRNA (in contrast to slow degradation of GFP) allows for more precise assessment of regulation of gene expression. To our surprise, although co-expression of *foxd3* and *ltk* is readily detected in 24-48 hpf larvae, overlap is seen in only around 50% of *ltk*-expressing cells at any of these stages. The criteria derived by our modelling with regard to the expression dynamics of factor R prompted us to conclude that *mitfa* repression cannot be fully explained by FoxD3 activity, although we cannot rule out this known transcriptional repressor (Kos et al. 2001; Ignatius et al. 2008; Thomas and Erickson 2009; Curran et al., 2009) making a partial contribution.

In summary, we have produced the first core GRN for the zebrafish iridophore, incorporating all known major players. This work now forms the basis for integration with our core GRN for the zebrafish melanocyte (Greenhill et al. 2011) in order to begin to see how integration of these GRNs is achieved in the pigment cell precursors enabling melanocyte versus iridophore fate choice. Whilst here we have focused on experimental genetics approaches and integrated mathematical modelling, a complementary approach looking at NC-specific histone marks would be informative, for example revealing active regulatory elements in iridophore lineage cells. However, a major priority will be to decode the mechanism repressing melanocyte (and potential other) fates in the iridophore lineage, with identification of factor R a crucial first step. Finally, continuous development of co-expression detection strategies (Lignell et al. 2017) will soon allow for simultaneously identifying an increasing number of marker genes, thus providing insight into the true potencies of partially restricted progenitors *in vivo*.

## Materials and methods

### Ethics statement

This study was performed with the approval of the University of Bath ethics committee and in full accordance with the Animals (Scientific Procedures) Act 1986, under Home Office Project Licenses 30/2937 and P87C67227.

### Fish husbandry

Embryos were obtained from natural crosses. Staging was performed according to Kimmel et al. (Kimmel et al. 1995). Unless stated otherwise, we used the WIK stock for experiments in WTs, and the following mutant lines: *sox10^t3^* (Dutton et al. 2001), *mitfa^w2^*(Lister et al. 1999), *ltk^ty82^* (Lopes et al. 2008) and *tfec^ba6^* (Petratou et al., 2018). Embryos were obtained by incrossing heterozygous carriers for each mutant allele, with WT siblings were used as controls.

### Transcript detection in whole mount embryos

Detailed information on the preparation of materials and the protocols for performing chromogenic whole mount in situ hybridisation (WISH) as well as multiplex fluorescent RNAscope can be found in Petratou et al. (Petratou et al. 2017; Petratou et al., in prep.). Probes used for chromogenic WISH were *sox10* (Dutton et al. 2001), *foxd3* (Odenthal and Nüsslein-Volhard 1998), *ltk* (Lopes et al. 2008), *pnp4a* (Curran et al. 2010), *mitfa* (Lister et al. 1999) and *tfec* (NM_001030105.2). To generate the *tfec* probe, cDNA prepared from total RNA extracted from 72 hpf zebrafish embryos was amplified with the following primers: forward 5’-AGCCAACAATCACGACAGTG-3’ and reverse 5’-CCAATAGAAACGGGAGGTCA-3’. The product was cloned into pCR II-BluntTOPO vector (Invitrogen) and the orientation assessed by sequencing. The plasmid was linearised with *PstI* restriction enzyme (NEB) and *in vitro* transcription was with the SP6 polymerase of the DIG labelling kit (Roche; Cat# 11175025910). For multiplex RNAscope, the following probes were used: *ltk* (ACD; Cat No. 444641), *tfec* (ACD; Cat No. 444701), *mitfa* (ACD; Cat No. 444651), *sox10* (ACD; Cat No. 444691) and *foxd3* (ACD; Cat No. 444681).

Embryos were imaged using an upright compound Imager 2 microscope (Zeiss). WISH samples were imaged under transmitted light, with an Axiocam 506 colour camera (Zeiss). RNAscope samples were imaged with dsRed, YFP and DAPI filters (supplied by Zeiss), using the Orca Flash 4.0 V2 camera (Zeiss) and Apotome.2 (Zeiss). Images were processed using the ZEN software (Zeiss), the FIJI package and Adobe Photoshop CS6.

We note that mutant and WT embryos were usually morphologically indistinguishable after fixation and were processed together. The Pearson’s chi-squared test (Pearson 1900; Harris 1912; Griffiths et al. 2000) was used to test the null hypothesis that in a sample of mixed WT, heterozygous and homozygous mutant embryos, observed alternative gene expression patterns correspond to the expected Mendelian ratios (75% of embryos are expected to show WT phenotype and 25% to potentially show altered gene expression). For this test, degrees of freedom = 1. The chi-squared table (Jones 2008) was used to calculate the probability that the number of observed embryos with an alternative expression phenotype was consistent with the expected number of homozygous mutants. For *p-value* > 0.1 the null hypothesis was accepted. For *p-value* < 0.1 it was assumed that alternative phenotypes in our samples were due to effects independent of the mutant genotype.

### Overexpression by microinjection

For overexpression assays, 50-70 pg of purified mRNA diluted in sterile water were injected in each WT (WIK) one-cell stage embryo using standard methods (Lister et al. 1999). Capped mRNA was prepared from plasmid templates using the SP6 mMessage mMachine kit (Ambion) for the overexpression of GFP, Sox10^WT^/Sox10^m618^ (Dutton et al. 2001) and Mitfa^WT^/Mitfa^b692^ (Lister et al. 1999; Lister, Close, and Raible 2001). For Tfec^WT^/Tfec^ba6^ overexpression, *in vitro* capped and polyadenylated mRNA was prepared using the mMessage mMachine T7 Ultra transcription kit (Ambion). Total RNA was isolated using TRI reagent (Sigma) from dissected trunks of 10-15 72 hpf WT (WIK) or homozygous *tfec^ba6^*^/ba6^ embryos and WT or mutant cDNA was generated using the SuperScript III First Strand Synthesis Supermix kit (Invitrogen). The *tfec* coding sequence (ENSDART00000164766.1) was amplified from the cDNA templates using the following primers: forward 5’-AGCGAGATCCTCCTGCTTCG-3’, reverse 5’-ATTCTGAGAGTGCGGTCCAG-3’. The T7 promoter was fused to the 5’ end of the amplicons through additional PCR amplification using the same reverse primer and the following forward: 5’- TAATACGACTCACTATAGGGAGAAGCGAGATCCTCCTGCTTCG-3’. The resulting amplicons were used as templates for *in vitro* transcription.

### Quantitative real-time PCR

Total RNA was extracted using TRI reagent (Sigma) from 8 embryos per sample at 6 hours post-injection. cDNA was synthesised from 1 μg of total RNA using the iScript cDNA synthesis kit (Biorad; Cat# 1708890). qRT-PCR was performed in duplicate using Fast SYBR Green Master Mix (Applied Biosystems; Cat# 4385617) and the StepOnePlus^TM^ Real-Time PCR System (Thermo Scientific; Cat# 4376600). For normalisation, we used expression of the housekeeping gene *rlp13* (primers ready-made from Primerdesign Ltd.). Primers for *tfec* transcript detection: forward 5’-GGAGCTTGGATTGCATGGAG-3’, reverse 5’-TTGATCAGCACCGTACACCT-3’. Primers for *pnp4a* transcript detection: forward 5’- TGGATGCAGTTGGAATGAGT-3’, reverse 5’-TTGACAGTCTCGTTGTCCTCA-3’. The ΔΔCt method (Livak and Schmittgen 2001) was used to evaluate relative changes in *pnp4a* expression, whereas absolute levels of *sox10* transcript were assessed using a standard curve. The null hypothesis that there was no change in the level of gene expression between control samples (injected with GFP or with null transcripts) and overexpression samples was rejected if *p-value* < 0.05 using a two-sample t-test without assuming equal variances. T-tests were performed using Microsoft Excel.

### Mathematical modelling

Modelling of gene interactions was done using the approach presented in Greenhill et al. (2011). For each model, gene expression dynamics were described using a system of ordinary differential equations (ODEs; see S1 Text). The equations were solved numerically using the *ode45* solver in MatLab software. Solving these equations returns the average concentration of gene output, measured in nM, across a homogeneous cell population. The Monte Carlo sampling algorithm used for randomising the constant parameters and subsequently scoring model outputs was run on MatLab. By random uniform logarithmic draw, we let all parameters vary in the range between a multiple 1/3.5 and 3.5. For each random draw, the system of ODEs of interest is solved and the resulting gene output dynamics are scored for biological relevance. For the scoring criteria refer to the results section, and for the corresponding mathematical functions see S2 Text.

## Acknowledgements

We gratefully acknowledge Dr G. Aquino and Dr F. Gubay for their advice and helpful comments regarding mathematical modelling and Dr R. D. Ballim and K. Camargo Sosa for technical advice. Work by JL was supported by National Institutes of Health Clinical and Translational Science Award (UL1TR000058 from the National Center for Advancing Translational Sciences) to Virginia Commonwealth University and the AD Williams’ Fund of Virginia Commonwealth University. Work by KP was supported by the University of Bath within the Biotechnology and Biological Sciences Research Council (BBSRC)-funded South West Biosciences Doctoral Training Partnership. Work by TS, RNK and HS was supported by BBSRC grant BB/L00769X/1 and by AR by BBSRC Grant BB/L007789/1.

## Supporting information

**S1 Table:** Statistics of loss of function experiments. The Pearson’s chi-squared test for goodness of fit indicates the likelihood that embryos presenting with an expression pattern differing from the WT might correspond to homozygous mutants of the respective allele (indicated in the second column). All the alleles follow the classic Mendelian ratios, thus 25% of the total of examined embryos are expected to be homozygous mutants. For each of the four stages, the 1^st^ sub-column presents the number of embryos out of the examined total which show a characteristic alternative expression pattern, and the 2^nd^ indicates the associated *p-value*. If p > 0.1 we accept the null hypothesis that any deviation in the observed number of embryos with an alternative phenotype from the expected number of homozygous mutants in the sample is only due to random chance. In red/bold are any samples where observed phenotypes did not significantly correlate with expected Mendelian ratios, i.e. where there is unlikely to be a mutant phenotype.

| WISH pattern | Genotype | Stages |  |  |  |  |  |  |  |
| --- | --- | --- | --- | --- | --- | --- | --- | --- | --- |
|  |  | 18 hpf |  | 24 hpf |  | 30 hpf |  | 48 hpf |  |
| tfec | sox10<sup>-/-</sup> | 0 of 41 | p < 0.001 | 20 of 74 | 0.6 < p < 0.7 | 17 of 66 | p ≈ 0.9 | 12 of 59 | 0.3 < p < 0.5 |
|  | ltk<sup>-/-</sup> | 0 of 79 | p < 0.001 | 5 of 194 | p < 0.001 | 23 of 80 | 0.3 < p < 0.5 | 29 of 99 | p ≈ 0.7 |
|  | tfec<sup>-/-</sup> |  |  | 14 of 48 | p ≈ 0.5 | 15 of 51 | 0.3 < p < 0.5 | 9 of 38 | 0.8 < p < 0.9 |
| ltk | tfec<sup>-/-</sup> |  |  | 12 of 49 | p ≈ 0.95 | 16 of 46 | 0.1 < p < 0.2 | 13 of 36 | 0.1 < p < 0.2 |
| pnp4a | sox10<sup>-/-</sup> |  |  | 13 of 45 | 0.5 < p < 0.7 | 8 of 39 | 0.4 < p < 0.5 | 11 of 33 | 0.2 < p < 0.3 |
|  | ltk<sup>-/-</sup> |  |  | 6 of 105 | p < 0.001 | 9 of 42 | 0.5 < p < 0.7 | 7 of 28 | p > 0.95 |
|  | tfec<sup>-/-</sup> |  |  | 11 of 49 | p ≈ 0.7 | 10 of 41 | p ≈ 0.95 | 9 of 44 | p ≈ 0.5 |
|  | mitfa<sup>-/-</sup> |  |  | 11 of 49 | 0.5 < p < 0.7 | 11 of 31 | 0.1 < p < 0.2 |  |  |

**S2 Table:**
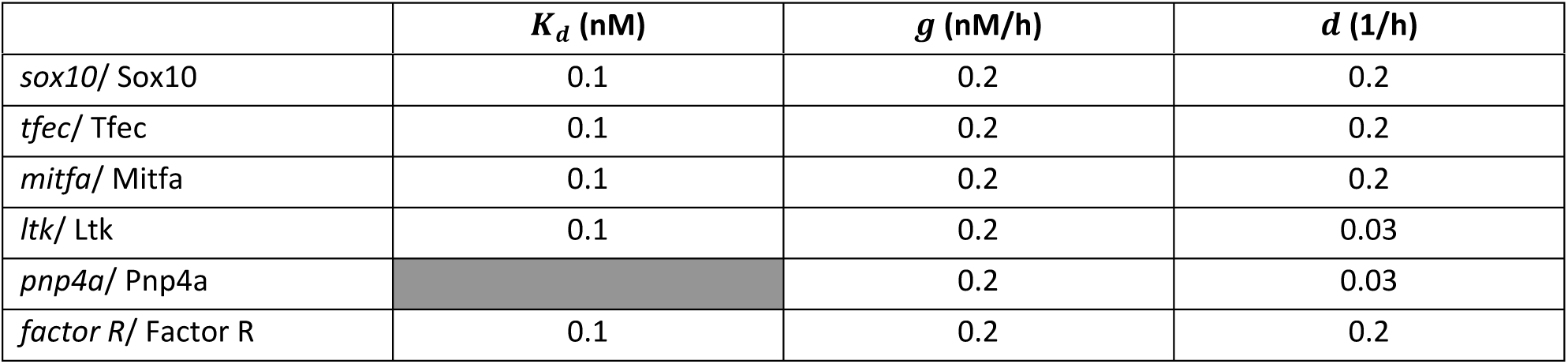
Parameter Choice. The default parameter set, selected as physiologically relevant based on published literature. These parameters were used for models A1, A2, A3 and B. For references see S1 Text.

| | $K_d$ (nM) | $g$ (nM/h) | $d$ (1/h) |
| --- | --- | --- | --- |
| *sox10/ Sox10* | 0.1 | 0.2 | 0.2 |
| *tfec/ Tfec* | 0.1 | 0.2 | 0.2 |
| *mitfa/ Mitfa* | 0.1 | 0.2 | 0.2 |
| *ltk/ Ltk* | 0.1 | 0.2 | 0.03 |
| *pnp4a/ Pnp4a* |  | 0.2 | 0.03 |
| *factor R/ Factor R* | 0.1 | 0.2 | 0.2 |

**S1 Figure.**
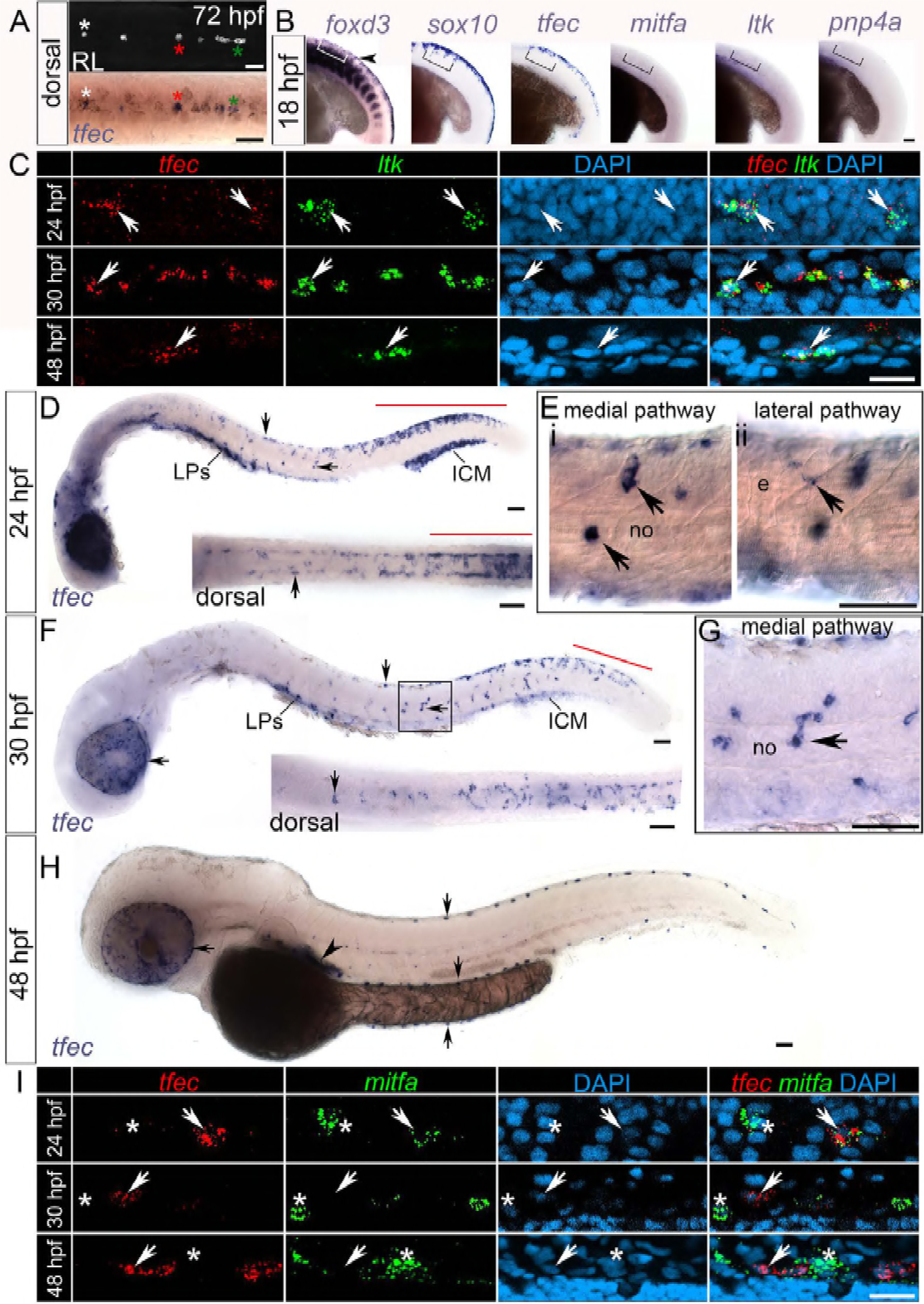
*pnp4a* expression is gradually reduced in presumptive iridoblasts, but not melanoblasts, in *ltk* mutants. Chromogenic WISH at 24 hpf shows almost complete elimination of *pnp4a* expression from the NC derivatives of the dorsal trunk (vertical arrowheads) and the migratory pathways (arrows) of *mitfa* mutants (B, B’), compared to WT siblings (A, A’). Expression in the RPE domain is reduced (horizontal arrowheads). WISH at 30 hpf (C-D’) reveals persistence of *pnp4a* expression in migrating cells which we interpret as melanoblasts, in *ltk* mutants (asterisks), as well as in the multipotent progenitor domain of the posterior tail (vertical arrowheads). We also observe a reduced number of cells in iridoblast locations: overlying the RPE (horizontal arrowheads), in the developing lateral patches and along the dorsal posterior trunk (arrows, enlarged in C’,D’). At 48 hpf (E,F), the majority of *pnp4a*- positive cells in iridophore locations are absent in *ltk* mutants. Specifically, cells overlying the RPE (horizontal arrowheads), on the lateral patches and along the dorsal and ventral posterior trunk and tail (arrows) are dramatically reduced. Very few escaper iridophores (F, arrows) maintain strong *pnp4a* expression upon loss of *ltk* function. LP, lateral patches. Lateral views, heads positioned towards the left. Scale bar corresponds to 100 μm in A,B,C,D,E,F and to 50 μm in A’,B’,C’,D’.

**S2 Figure.**
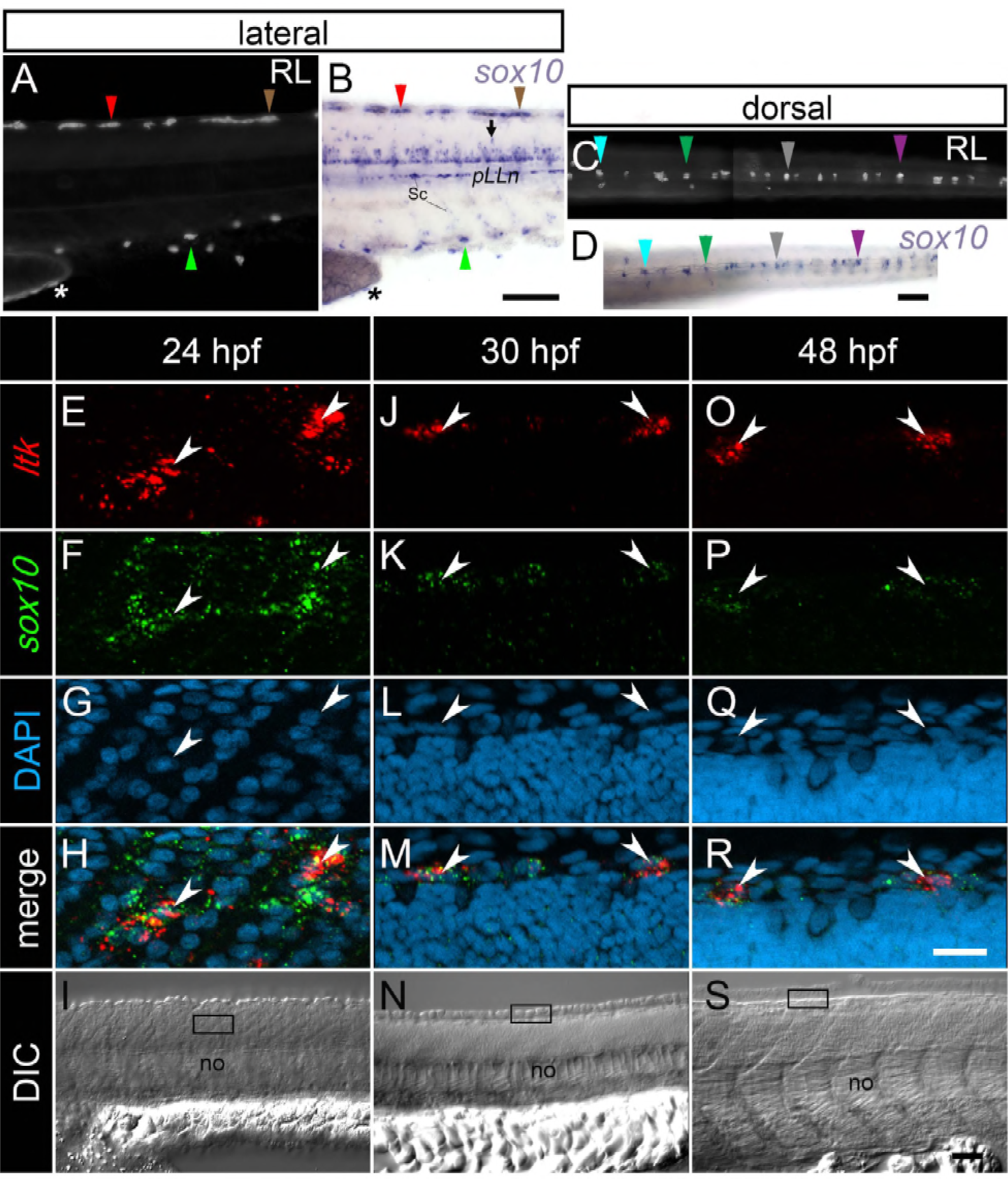
The mean cytoplasmic volume of cells in the dorsal ARPT can be calculated using high resolution DIC images combined with DAPI staining. (A) DIC image of a single focal plane from a Z-stack, showing the ARPT of a 24 hpf WT embryo. (B) Magnified view of the boxed region in (A). DIC allows for identification of the boundaries of the cells directly dorsal to the CNS (likely epidermal), while DAPI stain renders the nuclei visible. It is thus possible to measure the length (L) and width (W) of whole cells (arrowheads), as well as of their respective nuclei. CNS, central nervous system; no, notochord; YSE, yolk sac extension. Lateral view, head positioned towards the left. Scale bar: A: 50 μm; B: 20 μm.

**S3 Figure.**
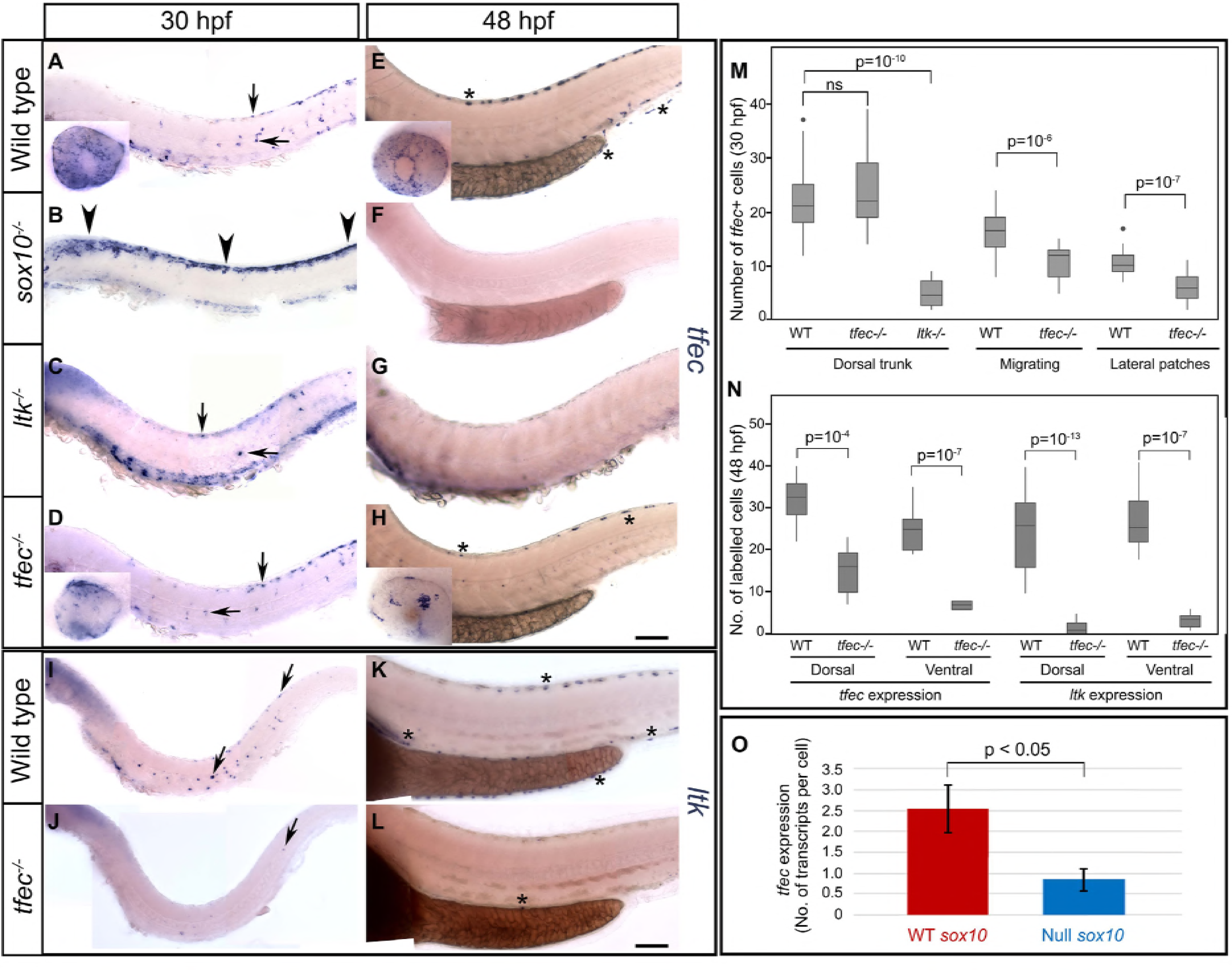
*In silico* testing of alternative Model B variants suggested that the currently included transcriptional regulators are insufficient for *mitfa* repression. Testing alternative methods of repressing *mitfa* expression using the mathematical model failed to recapitulate the experimentally observed *mitfa* dynamics in the absence of factor R. (A,B) Implementing Tfec-dependent suppression of *mitfa* resulted only in a relatively lower positive plateau of Mitfa output, instead of a peak at approximately 24 hpf, followed by downregulation of *mitfa*. (C,D) Implementing Sox10-dependent suppression of *mitfa* resulted in the same outcome.

**S4 Figure.**
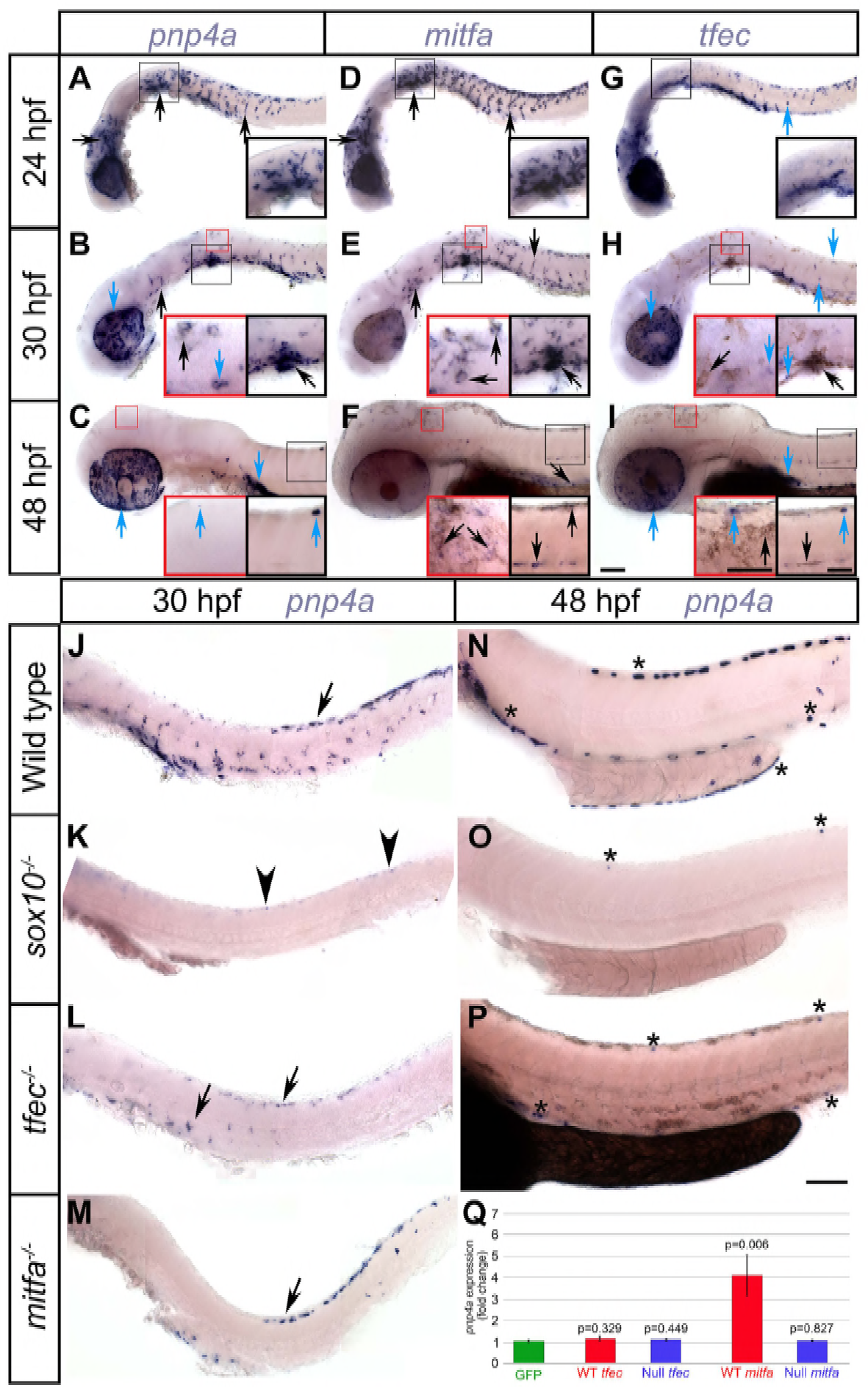
*foxd3* is not a good candidate for the role of factor R in the iridophore lineage. RNAscope experiments at 36 hpf reveal that *foxd3* does not fulfil the criteria established using our models for factor R, as it is only expressed in a subset (approximately 50%) of *ltk*+ Ib(df) of the posterior dorsal trunk. In (B) arrowheads point at *ltk*+ cells that co-express *foxd3*, while asterisks indicate cells that are only positive for *ltk*. no, notochord; YSS, yolk sac stripe. Lateral views, head towards the left. Scale bars: 50 μm.

**S5 Figure.**
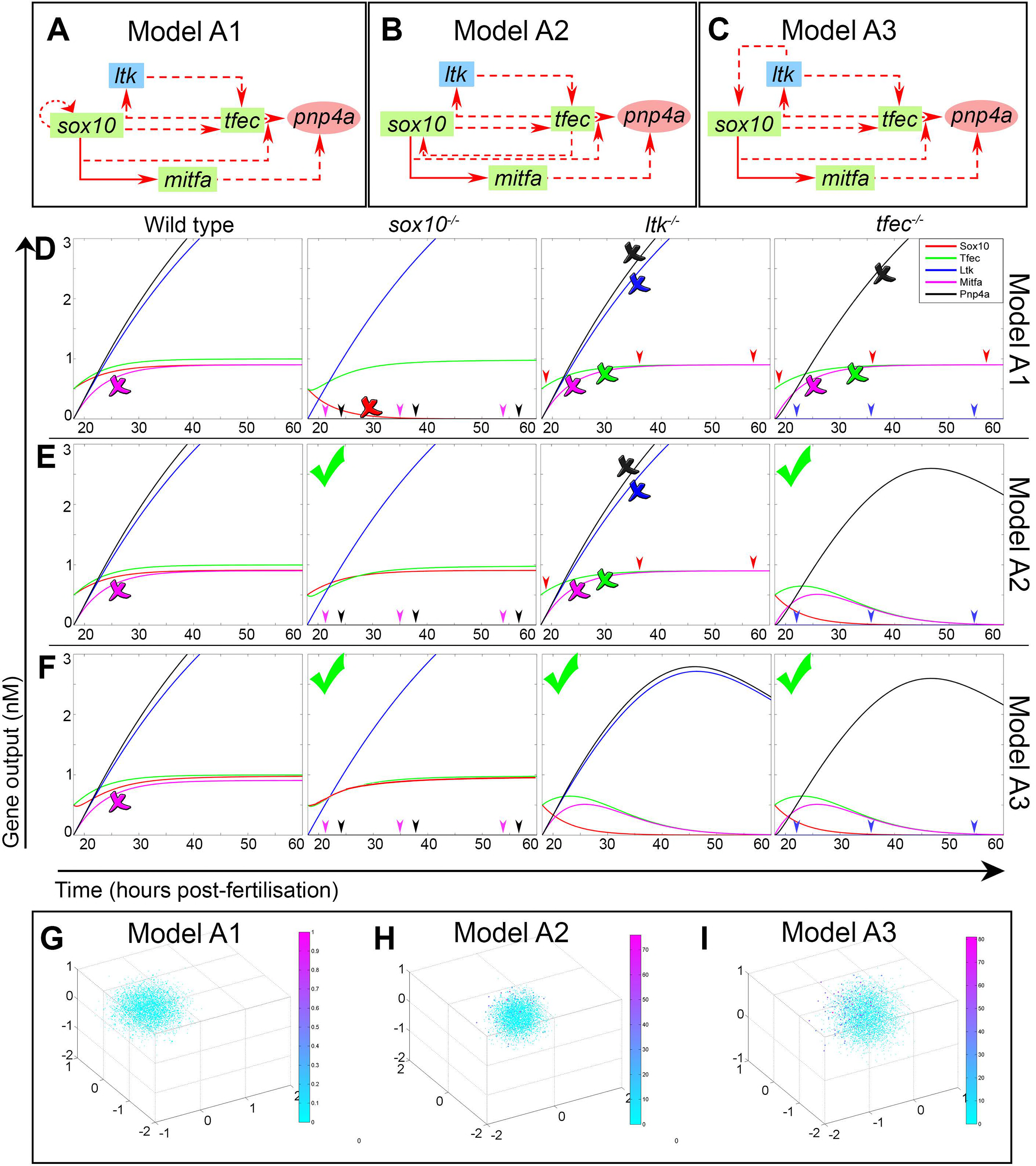
Best achievable outputs using Monte Carlo regarding *mitfa* expression dynamics under all of our models. WT outputs derived from the three best scoring parameter combinations (i, ii, iii) following 20,000 runs using Monte Carlo for models A1 (A), A2 (B), A3 (C), B(2) (D) and B(3) (E; referred to as model B in this work). (i’, ii’ and iii’) represent magnified *mitfa* expression dynamics in each of the outputs. In model A1 (Ai-Aiii’), a very subtle rise and drop of Mitfa concentration is only achievable if Sox10 concentration reaches undetectable levels at steady-state (red arrowheads). In (Bi) Mitfa remains very low throughout iridoblast specification ([M]<0.1 nM), while in (Bii), (Biii) the decline is very subtle and mitfa remains upregulated in steady state. (Ci) and (Ciii) show Mitfa staying relatively high at steady state, compared to the achieved maximum levels, with Sox10 declining below detection level in (Ciii). In (Cii) Mitfa is steadily upregulated to its steady-state concentration value. In the two highest scoring model B(2) outputs (where factor R was incorporated in model A2), Mitfa exhibits the required rise and drop dynamics only when Sox10 declines (Di, Dii; red arrowheads). The WT outputs of the three top scoring trials for model B(3) (derived from A3), exhibit maintenance of sox10 expression, while Mitfa peaks at approximately 24 hpf, before declining to no more than half the maximum value (Ei-Eiii’). PCA analysis of all Monte Carlo outputs for this model (F) reveals that significantly fewer trials score high (dark blue and magenta spots) and that the absolute score value is lower, compared to trials using model B(3), i.e. the chosen model B (G).

**S1 Text Derivation of the system of ODEs and assignment of parameter constants.**

**S2 Text Monte Carlo scoring functions.**

## References

Adameyko, I., and F. Lallemend. 2010. “Glial versus Melanocyte Cell Fate Choice: Schwann Cell Precursors as a Cellular Origin of Melanocytes.” Cellular and Molecular Life Sciences.

Adameyko, I., F. Lallemend, J. B. Aquino, J. Pereira, P. Topilko, T. Müller, N. Fritz, et al. 2009. “Schwann Cell Precursors from Nerve Innervation Are a Cellular Origin of Melanocytes in Skin.” Cell 139 (2). Elsevier Ltd: 366–79.

Bagnara, J. T., J. Matsumoto, W. Ferris, S. K. Frost, W. A. Turner, T. T. Tchen, and J. D. Taylor. 1979. “Common Origin of Pigment Cells.” Science 203 (4379): 410–15.

Bagnara, J.T. 2007. “Comparative Anatomy and Physiology of Pigment Cells in Nonmammalian Tissues.” In The Pigmentary System Physiology and Pathophysiology, edited by JJ Nordland, RE Boissy, VJ Hearing, RA King, and JP Ortonne, 9–40. New York: Oxford University Press.

Curran, K., J. A. Lister, G. R. Kunkel, A. Prendergast, D. M. Parichy, and D. W. Raible. 2010. “Interplay between Foxd3 and Mitf Regulates Cell Fate Plasticity in the Zebrafish Neural Crest.” Developmental Biology 344 (1). Elsevier Inc.: 107–18.

Curran, K., D. W. Raible, and J. A. Lister. 2009. “Foxd3 Controls Melanophore Specification in the Zebrafish Neural Crest by Regulation of Mitf.” Developmental Biology 332 (2). Elsevier Inc.: 408–17.

Douarin, N. M. Le, G. W. Calloni, and E. Dupin. 2008. “The Stem Cells of the Neural Crest.” Cell Cycle 7 (8). Landes Bioscience: 1013–19.

Douarin, N. M. Le, and E. Dupin. 2003. “Multipotentiality of the Neural Crest.” Current Opinion in Genetics & Development 13 (5): 529–36.

Dutton, K. A., A. Pauliny, S. S. Lopes, S. Elworthy, T. J. Carney, J. Rauch, R. Geisler, P. Haffter, and R. N. Kelsh. 2001. “Zebrafish Colourless Encodes sox10 and Specifies Non-Ectomesenchymal Neural Crest Fates.” Development 128 (21): 4113–25.

Elworthy, S., J. A. Lister, T. J. Carney, D. W. Raible, and R. N. Kelsh. 2003. “Transcriptional Regulation of Mitfa Accounts for the sox10 Requirement in Zebrafish Melanophore Development.” Development 130 (12): 2809–18.

Fadeev, A., J. Krauss, A. P. Singh, and C. Nüsslein-Volhard. 2016. “Zebrafish Leucocyte Tyrosine Kinase Controls Iridophore Establishment, Proliferation and Survival.” Pigment Cell & Melanoma Research 29 (3): 284–96.

Gilbert, S. F. 2000. “The Neural Crest.” In Developmental Biology, 6th ed. Sunderland (MA): Sinauer Associates.

Greenhill, E. R., A. Rocco, L. Vibert, M. Nikaido, and R. N. Kelsh. 2011. “An Iterative Genetic and Dynamical Modelling Approach Identifies Novel Features of the Gene Regulatory Network Underlying Melanocyte Development.” Edited by Mary C Mullins. PLoS Genetics 7 (9). Public Library of Science: 18.

Griffiths, A. J. F., J. H. Miller, D. T. Suzuki, R. C. Lewontin, and W. M. Gelbart. 2000. “Chi-Square Test.” W. H. Freeman.

Harris, A. J. 1912. “A Simple Test of the Goodness of Fit of Mendelian Ratios.” The American Naturalist 46 (552): 741–45.

Higdon, C. W., R. D. Mitra, and S. L. Johnson. 2013. “Gene Expression Analysis of Zebrafish Melanocytes, Iridophores, and Retinal Pigmented Epithelium Reveals Indicators of Biological Function and Developmental Origin.” PLoS One 8 (7): e67801.

Ignatius, M. S., H. E. Moose, H. M. El-Hodiri, and P. D. Henion. 2008. “colgate/hdac1 Repression of foxd3 Expression Is Required to Permit Mitfa-Dependent Melanogenesis.” Developmental Biology 313 (2): 568–83.

Jones, J. 2008. “Table: Chi-Square Probabilities.” http://people.richland.edu/james/lecture/m170/tbl-chi.html.

Kelsh, R. N. 2004. “Genetics and Evolution of Pigment Patterns in Fish.” Pigment Cell & Melanoma Research 17 (4): 326–36.

Kelsh, R. N., M. Brand, Y. J. Jiang, C. P. Heisenberg, S. Lin, P. Haffter, J. Odenthal, et al. 1996. “Zebrafish Pigmentation Mutations and the Processes of Neural Crest Development.” Development 123 (1): 369–89.

Kelsh, R. N., M. L. Harris, S. Colanesi, and C. A. Erickson. 2009. “Stripes and Belly-Spots -- a Review of Pigment Cell Morphogenesis in Vertebrates.” Seminars in Cell Developmental Biology 20 (1). Elsevier Ltd: 90–104.

Kelsh, R.N. 2006. “Sorting out Sox10 Functions in Neural Crest Development.” BioEssays News and Reviews in Molecular Cellular and Developmental Biology 28 (8). Wiley Subscription Services, Inc., A Wiley Company: 788–98.

Kim, J., L. Lo, E. Dormand, and D. J. Anderson. 2003. “SOX10 Maintains Multipotency and Inhibits Neuronal Differentiation of Neural Crest Stem Cells.” Neuron 38 (1): 17–31.

Kimmel, C. B., W. W. Ballard, S. R. Kimmel, B. Ullmann, and T. F. Schilling. 1995. “Stages of Embryonic Development of the Zebrafish.” Developmental Dynamics 203 (3). Wiley Online Library: 253–310.

Kimura, T., Y. Takehana, and K. Naruse. 2017. “pnp4a Is the Causal Gene of the Medaka Iridophore Mutantguanineless.” G3 (Bethesda, Md.) 7 (4). Genetics Society of America: 1357–63.

Kléber, M., H.-Y. Lee, H. Wurdak, J. Buchstaller, M. M. Riccomagno, L. M. Ittner, U. Suter, D. J. Epstein, and L. Sommer. 2005. “Neural Crest Stem Cell Maintenance by Combinatorial Wnt and BMP Signaling.” The Journal of Cell Biology 169 (2). The Rockefeller University Press: 309–20.

Kos, R., M. V. Reedy, R. L. Johnson, and C. A. Erickson. 2001. “The Winged-Helix Transcription Factor FoxD3 Is Important for Establishing the Neural Crest Lineage and Repressing Melanogenesis in Avian Embryos.” Development 128 (8): 1467–79.

Lignell, A., L. Kerosuo, S. J. Streichan, L. Cai, and M. E. Bronner. 2017. “Identification of a Neural Crest Stem Cell Niche by Spatial Genomic Analysis.” Nature Communications 8 (1). Nature Publishing Group: 1830.

Lister, J. A., J. Close, and D. W. Raible. 2001. “Duplicate Mitf Genes in Zebrafish: Complementary Expression and Conservation of Melanogenic Potential.” Developmental Biology 237 (2): 333–44.

Lister, J. A., C. Cooper, K. Nguyen, M. Modrell, K. Grant, and D. W. Raible. 2006. “Zebrafish Foxd3 Is Required for Development of a Subset of Neural Crest Derivatives.” Developmental Biology 159 (1). Elsevier Inc.: 50–59.

Lister, J. A., B. M. Lane, A. Nguyen, and K. Lunney. 2011. “Embryonic Expression of Zebrafish MiT Family Genes tfe3b, Tfeb, and Tfec.” Developmental Dynamics 240 (11): 2529–38.

Lister, J. A., C. P. Robertson, T. Lepage, S. L. Johnson, and D. W. Raible. 1999. “Nacre Encodes a Zebrafish Microphthalmia-Related Protein That Regulates Neural-Crest-Derived Pigment Cell Fate.” Development 126 (17): 3757–67.

Livak, K. J., and T. D. Schmittgen. 2001. “Analysis of Relative Gene Expression Data Using Real-Time Quantitative PCR and the 2−∆∆CT Method.” Methods 25 (4): 402–8.

Lopes, S. S., X. Yang, J. Müller, T. J. Carney, A. R. McAdow, G.-J. Rauch, A. S. Jacoby, et al. 2008. “Leukocyte Tyrosine Kinase Functions in Pigment Cell Development.” Edited by Greg Barsh. PLoS Genetics 4 (3). Public Library of Science: 13.

Minchin, J. E. N., and S. M. Hughes. 2008. “Sequential Actions of Pax3 and Pax7 Drive Xanthophore Development in Zebrafish Neural Crest.” Developmental Biology 317 (2): 508–22.

Nagao, Y., T. Suzuki, A. Shimizu, T. Kimura, R. Seki, T. Adachi, C. Inoue, et al. 2014. “Sox5 Functions as a Fate Switch in Medaka Pigment Cell Development.” Edited by David M. Parichy. PLoS Genetics 10 (4): e1004246.

Nagao, Y., H. Takada, M. Miyadai, T. Adachi, R. Seki, Y. Kamei, I. Hara, et al. 2018. “Distinct Interactions of Sox5 and Sox10 in Fate Specification of Pigment Cells in Medaka and Zebrafish.” PLoS Genetics.

Odenthal, J., and C. Nüsslein-Volhard. 1998. “Fork Head Domain Genes in Zebrafish.” Development Genes and Evolution 208 (5): 245–58.

Paratore, C., C. Eichenberger, U. Suter, and L. Sommer. 2002. “Sox10 Haploinsufficiency Affects Maintenance of Progenitor Cells in a Mouse Model of Hirschsprung Disease.” Human Molecular Genetics 11 (24): 3075–85.

Parichy, D. M., E. M. Mellgren, J. F. Rawls, S. S. Lopes, R. N. Kelsh, and S. L. Johnson. 2010. “Mutational Analysis of Endothelin Receptor b1 (Rose) during Neural Crest and Pigment Pattern Development in the Zebrafish Danio Rerio.” Developmental Biology 227 (2): 294–306.

Pearson, K. 1900. “X. On the Criterion That a given System of Deviations from the Probable in the Case of a Correlated System of Variables Is Such That It Can Be Reasonably Supposed to Have Arisen from Random Sampling.” Philosophical Magazine Series 5 50 (302). Taylor & Francis: 157–75.

Petratou, K., K. C. Sosa, R. Al Jabri, Y. Nagao, and R. N. Kelsh. 2017. “Neural Crest Methodologies in Zebrafish and Medaka: Transcript and Protein Detection Methodologies for Neural Crest Research on Whole Mount Zebrafish and Medaka.” Springer.

Potterf, S. B., M. Furumura, K. J. Dunn, H. Arnheiter, and W. J. Pavan. 2000. “Transcription Factor Hierarchy in Waardenburg Syndrome: Regulation of MITF Expression by SOX10 and PAX3.” Human Genetics 107 (1): 1–6.

Raible, D. W., and J. S. Eisen. 1994. “Restriction of Neural Crest Cell Fate in the Trunk of the Embryonic Zebrafish.” Development 120: 495–503.

Rodrigues, F. S. L. M., X. Yang, M. Nikaido, Q. Liu, and R. N. Kelsh. 2012. “A Simple, Highly Visual in Vivo Screen for Anaplastic Lymphoma Kinase Inhibitors.” ACS Chemical Biology 7 (12): 1968–74.

Schartl, M., L. Larue, M. Goda, M. W. Bosenberg, H. Hashimoto, and R. N. Kelsh. 2016. “What Is a Vertebrate Pigment Cell?” Pigment Cell & Melanoma Research 29 (1): 8–14.

Shakhova, O., and L. Sommer. 2008. Neural Crest-Derived Stem Cells. StemBook. Harvard Stem Cell Institute.

Thomas, A. J., and C. A. Erickson. 2009. “FOXD3 Regulates the Lineage Switch between Neural Crest-Derived Glial Cells and Pigment Cells by Repressing MITF through a Non-Canonical Mechanism.” Development 136 (11). Company of Biologists: 1849–58.

Trainor, P. A. 2014. Neural Crest Cells: Evolution, Development and Disease. Elsevier/AP.

Vibert, L., G. Aquino, I. Gehring, T. Subkankulova, T. F. Schilling, A. Rocco, and R. N. Kelsh. 2017. “An Ongoing Role for Wnt Signaling in Differentiating Melanocytes in Vivo.” Pigment Cell & Melanoma Research 30 (2): 219–32.

Waddington, C. H. 1957. The Strategy of the Genes: A Discussion of Some Aspects of Theoretical Biology. London: George Allen & Unwin.

Weston, J. A. 1991. “Sequential Segregation and Fate of Developmentally Restricted Intermediate Cell Populations in the Neural Crest Lineage.” Current Topics in Developmental Biology 25 (January): 133–53.

